# On stability of Canonical Correlation Analysis and Partial Least Squares with application to brain-behavior associations

**DOI:** 10.1101/2020.08.25.265546

**Authors:** Markus Helmer, Shaun Warrington, Ali-Reza Mohammadi-Nejad, Jie Lisa Ji, Amber Howell, Benjamin Rosand, Alan Anticevic, Stamatios N. Sotiropoulos, John D. Murray

**Author notes:** (JDM), (SNS).

## Abstract

Associations between datasets can be discovered through multivariate methods like Canonical Correlation Analysis (CCA) or Partial Least Squares (PLS). A requisite property for interpretability and generalizability of CCA/PLS solutions is stability of feature patterns driving an association. However, stability of CCA/PLS in high-dimensional datasets is questionable, as found in empirical characterizations. To study these issues in a systematic manner, we developed a generative modeling framework to simulate synthetic datasets, parameterized by dimensionality, variance structure, and association strength. We found that when sample size is relatively small, but comparable to typical studies, CCA/PLS associations are highly unstable and inaccurate; both in their magnitude and importantly in the latent pattern underlying the discovered association. We confirmed these trends across two neuroimaging modalities, functional and diffusion MRI, and in independent datasets, Human Connectome Project (n*≈*1000) and UK Biobank (n*≈*20000) and found that only the latter comprised sufficient samples for stable mappings between imaging-derived and behavioral features. We further developed a power calculator to provide sample sizes required for stability and reliability of multivariate analyses for future studies.

## Introduction

Discovery of associations between high-dimensional datasets is a topic of growing importance across scientific disciplines. For instance, large initiatives in human neuroimaging collect, across thousands of subjects, rich multivariate neural measures paired with psychometric and demographic measures ^1, 2^. A major goal is to determine the existence of an association linking individual variation in behavioral features to variation in neural features and to characterize the dominant latent patterns of features that underlie this association ^3, 4^. One widely employed statistical approach to map multivariate associations is to define linearly weighted composites of features in both datasets (e.g., neural and psychometric) with the sets of weights—which correspond to axes of variation—selected to maximize between-dataset association strength (Fig. 1A). The resulting profiles of weights for each dataset can be examined for how the features form the association. Depending on whether association strength is measured by correlation or covariance, the method is called *canonical correlation analysis* (CCA) ^5^ or *partial least squares* (PLS) ^6–11^, respectively. CCA and PLS are commonly employed across scientific fields, including genomics ^12^ and neuroimaging ^3, 4, 13, 14^.

**Figure 1.**
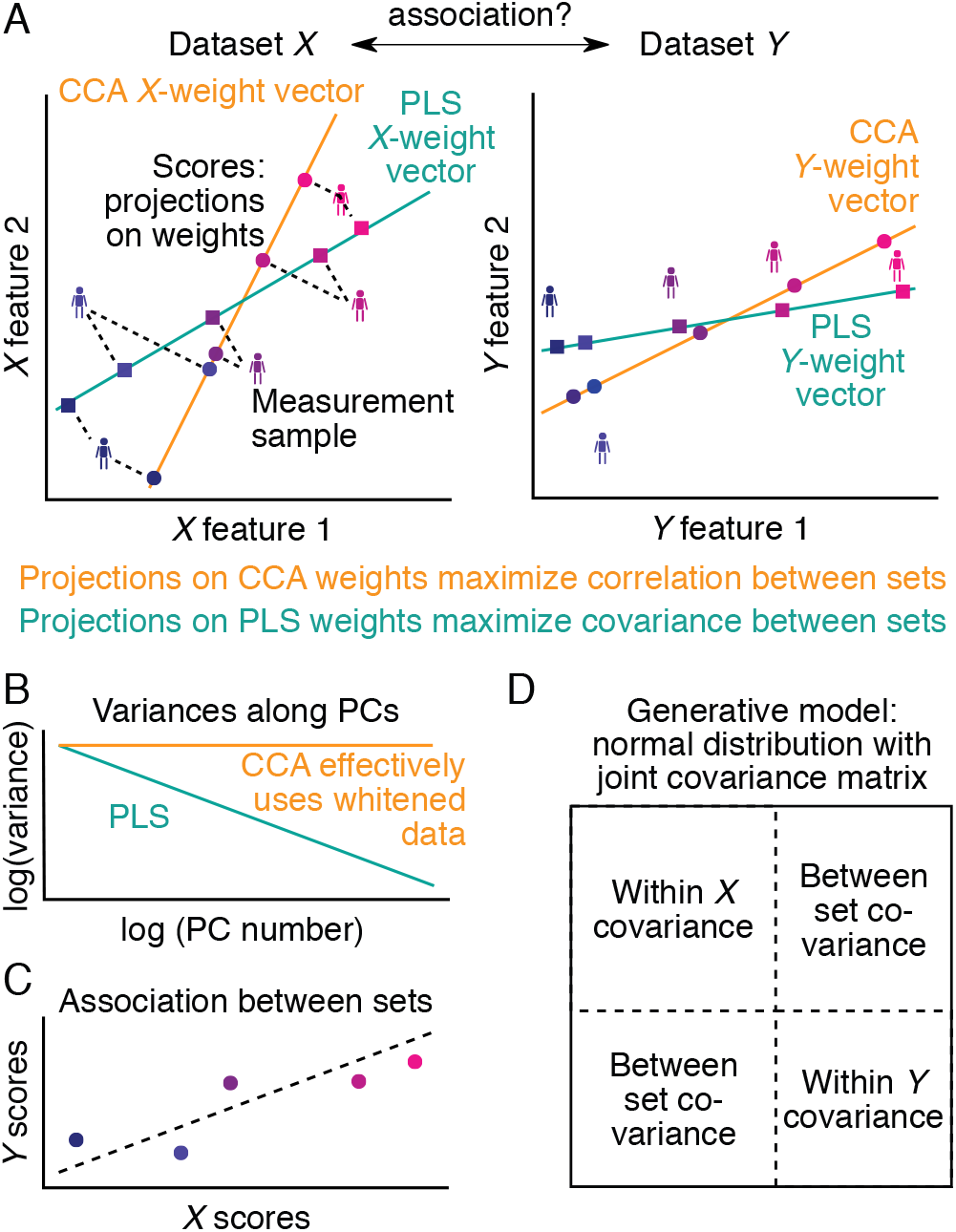
Overview of CCA, PLS and the generative model used to investigate their properties. **A)** Two multivariate datasets, *X* and *Y*, are projected separately onto respective weight vectors, resulting in univariate scores for each dataset. The weight vectors are chosen such that the correlation (for CCA) or covariance (for PLS) between *X* and *Y* scores is maximized. **B)** In the principal component coordinate system, the variance structure within each dataset can be summarized by its principal component spectrum. For simplicity, we assume that these spectra can be modeled as power-laws. CCA, uncovering correlations, disregards the variance structure and can be seen as effectively using whitened data (cf. Methods). **C)** The association between sets is encoded in the association strength of *X* and *Y* scores. **D)** Datasets *X* and *Y* are jointly modeled as a multivariate normal distribution. The within-set variance structure (**B**) corresponds to the blocks on the diagonal, and the associations between datasets (**C**) are encoded in the off-diagonal blocks.

Analysis of such high-dimensional datasets is challenging due to inherent measurement noise and the often small sample sizes in comparison to the dimensionality of the data. Although the utility of CCA/PLS is well established, open challenges exist regarding stability in characteristic regimes of dataset properties. Stability implies that elements of CCA/PLS solutions, such as association strength and weight profiles, are reliably estimated across independent samples from the same population. Instability or overfitting can occur if an insufficient sample size is available to properly constrain the model. Manifestations of instability and overfitting in CCA/PLS include inflated association strengths ^15–19^, out-of-sample association strengths markedly lower than in-sample ^18, 20^, and feature profiles/patterns that vary substantially across studies ^15, 18–25^. Stability of models is essential for replicability, generalizability, and interpretability ^26^. Therefore, it is important to understand how stability of CCA/PLS solutions depends on dataset properties.

In neuroimaging, it has recently been suggested that thousands of subjects are required to achieve reproducible results when performing multivariate “brain-wide association studies,” as effect sizes are typically small ^27^. This claim generated recent debate in the field ^28–31^. A number of papers argue that larger effect sizes can be expected ^28, 29^, that sample-size requirements can be reduced via focused designs and cohorts ^30^, and that cross-validation avoids inflated associations ^31^. Yet, all previous studies and comments are mostly based on empirical observations and focus primarily on effect sizes. In the context of CCA/PLS, it remains unclear how elements of solutions differentially depend on dataset properties, and how CCA vs. PLS as distinct methods exhibit differential robustness across dataset regimes.

To investigate these issues systematically and go beyond empirical observations, we developed a generative statistical model to simulate synthetic datasets with known latent axes of association. Sampling from the generative model allows quantification of deviations between estimated and true CCA/PLS solutions. We found that stability of CCA/PLS solutions requires more samples (per feature) than are commonly used in published neuroimaging studies. With too few samples, estimated association strengths were too high, and estimated weights could be unreliable and non-generalisable for interpretation. CCA and PLS differed in their dependencies and robustness, in part due to PLS weights exhibiting an increased similarity towards dominant principal component axes compared to CCA weights. We analyzed two large state-of-the-art neuroimaging-psychometric datasets, the Human Connectome Project ^1^ and the UK Biobank ^2^, which followed similar trends as our model. We also observed similar trends when considering features from two neuroimaging modalities, functional and diffusion MRI. These model and empirical findings, in conjunction with a meta-analysis of estimated stability in the brain-behavior CCA literature, suggest that discovered association patterns through typical CCA/PLS studies in neuroimaging are prone to instability. Finally, we applied the generative model to develop algorithms and a software package for calculation of estimation errors and required sample sizes for CCA/PLS. We end with practical recommendations for application and interpretation of CCA/PLS in future studies.

## Results

CCA/PLS describe statistical associations between multivariate datasets by analyzing their between-set covariance matrix (Fig. 1A). A weighted combination of features (“scores”) is formed for each of the two datasets, and the association strength between these score vectors is optimized by defining the “weight” vectors. CCA and PLS use Pearson correlation and covariance as their objective functions, respectively. (PLS is also referred to as PLS correlation [PLSC] or PLS-SVD ^6–11^.) We call the corresponding optimized value “between-set correlation” (or “canonical correlation”) and “between-set covariance”, respectively. We also calculate “loadings”, which we define as the univariate Pearson correlations (across observations) between CCA/PLS scores and each original variable in the dataset. These terminologies are not used consistently across the literature (see Methods) ^4, 8, 13, 32, 33^.

### A generative model for cross-dataset multivariate associations

To analyze dependencies of stability for CCA and PLS, we need to generate synthetic datasets of stochastic samples with known, controlled properties. We therefore developed a generative statistical modeling framework, GEMMR (Generative Modeling of Multivariate Relationships), which allows us to design and generate synthetic datasets, investigate the dependence of CCA/PLS solutions on dataset size and assumed covariances, estimate weight errors in CCAs reported in the literature, and calculate sample sizes required to bound estimation errors (see Methods).

To describe GEMMR, first note that data for CCA/PLS consist of two datasets, given as data matrices *X* and *Y*, each with multiple features and an equal number *n* of samples. We model the within-set covariance with power-law decay in the variance spectrum, which we constrain to empirically consistent ranges (Fig. S1). GEMMR then embeds between-set associations by defining associated weight axes in each set. Finally, the joint covariance matrix for *X* and *Y* is composed using the within- and between-set covariances (Fig. 1D) and the normal distribution associated with this joint covariance matrix constitutes our generative model.

We systematically investigated the downstream effects on CCA/PLS stability of generative model parameters for dataset properties: number of features, assumed population (or true) value of between-set correlation, power-laws describing the within-set variances, and sample size. Weight vectors were chosen randomly and constrained such that the *X* and *Y* scores explain at least half as much variance as an average principal component in their respective sets. For simplicity, we restrict our present analyses to a single between-set association mode. We use “number of features” to denote the total number across both *X* and *Y*.

### Sample-size dependence of estimation error

Using surrogate datasets from our generative model, we characterized estimation error in multiple elements of CCA/PLS solutions. Here we use “samples per feature” as an effective sample-size measurement, which accounts for widely varying dimensionalities across empirical datasets (Fig. S2). A typical sample size in the brain-behavior CCA/PLS literature is about 5 samples per feature (Fig. S3A), marked with a dashed vertical line in Fig. 2. A key parameter of the generative model is the population value, or “true” value, of the association strength, i.e., the value one would obtain, both through in-sample and out-of-sample estimation, as the sample size grows toward infinity. Importantly, like the mean of a normal distribution the population value of the association strength, *r*_true_, is independent of the sample and the sample size used to estimate it, but constitutes instead a parameter of the distribution from which samples are drawn. As such *r*_true_ is a well-defined free parameter that can be varied independently of sample size.

**Figure 2.**
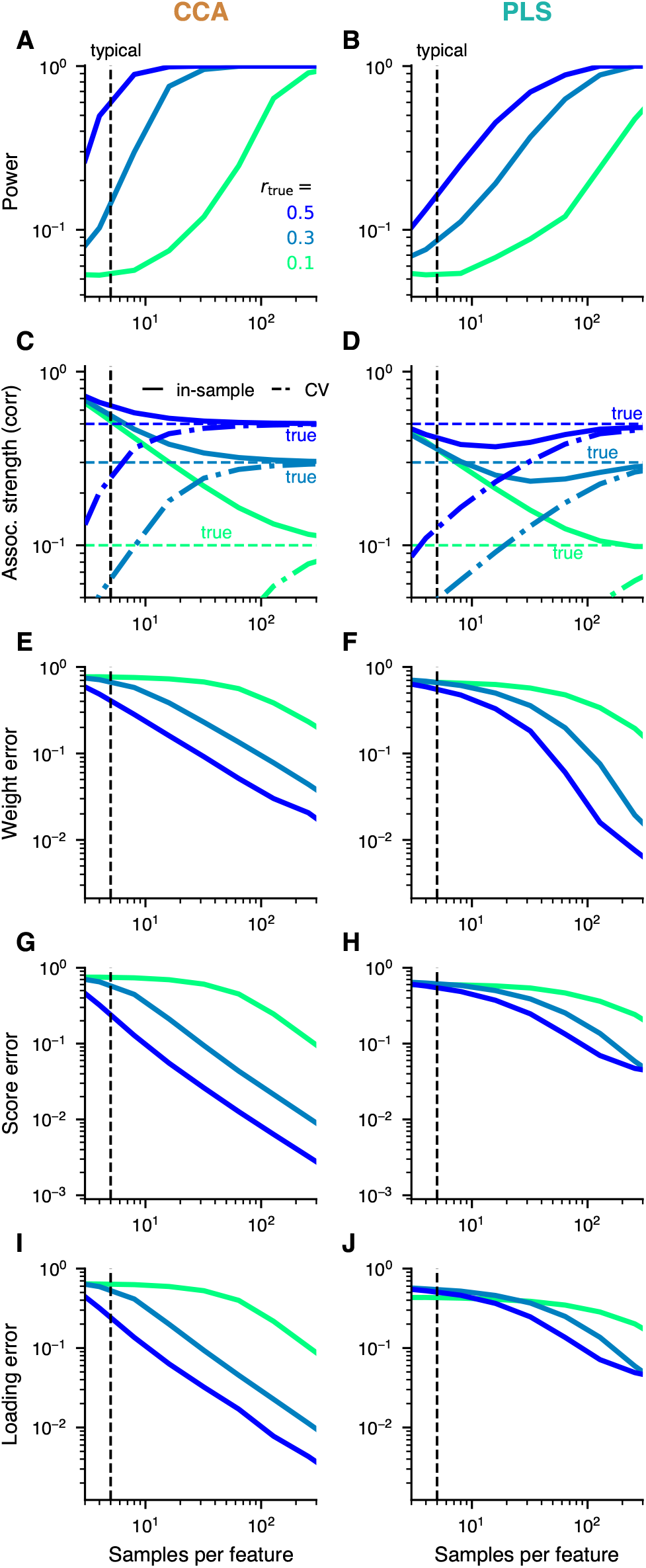
Sample-size dependence of CCA and PLS. **(A, B)** For sufficiently large sample sizes, statistical power to detect a non-zero correlation converges to 1. **(C, D)** In-sample (solid) and cross-validated (dash-dotted) estimates of the between-set correlations approach their assumed true (population) value (dashed). **(E, F)** Weight errors (quantified as the absolute cosine distance between the true weights of the generative model and estimated weights from CCA/PLS on the sample, separately for *X* and *Y* and taking the greater of the two), **(G, H)** score errors (measured as “1 - Pearson correlation” between estimated and true scores, which, in turn, are obtained by applying estimated and true weights to common test data) **(I, J)** as well as loading errors (measured as “1 - Pearson correlation” between estimated and true loadings) become close to 0. Original data features are generally different from principal component scores, but as the relation between these two data representations cannot be constrained, we calculate all loadings here with respect to principal component scores. Moreover, to compare loadings across repeated datasets we calculate loadings for a common test set, as for CCA/PLS scores. Left and right columns show results for CCA and PLS, respectively. For all metrics, convergence depends on the true (population) between-set correlation *r*_true_ and is slower if *r*_true_ is low. Note that the color code indicates true (population) between-set correlation and corresponds to the dashed horizontal lines in C-D. The dashed vertical line at 5 samples per feature represents a typically used value (Fig. S3A). All curves in this figure are averaged across samples obtained from a number of different generative models with varying true (population) weight vectors and dimensionality (see Methods).

We assessed, first, whether a significant association can robustly be detected, quantified by statistical power, and found relatively low power at typical sample sizes and population effect sizes (Fig. 2A-B). Second, we evaluated convergence of association strength (Fig. 2C-D). We calculated the “(with-)in sample” association strength by performing CCA/PLS with a given sample of data, and “out-of-sample” association strength through cross-validation ^20^. In-sample estimates of the association strength overestimate their true value (Figs. 2C, S4, S5). A sufficient sample size, depending on other covariance matrix properties, is needed to bound the error in the association strength. The observed correlation converges to its assumed true (population) value for sufficiently large sample sizes (Fig. 2C-D). Cross-validated estimates underestimate the true value to a similar degree as in-sample estimates overestimate it ^18^ (Fig. S6).

In addition to association strengths, CCA/PLS solutions provide weights that encode the nature of the association in each dataset, as well as scores which represent a latent value assigned to each sample (e.g., subject). Finally, some studies report loadings, i. e. the correlations between original data features and CCA/PLS scores (Fig. S7A-B). We found that estimation errors for weights, scores, and loadings decrease monotonically with sample size and more quickly for stronger population effect sizes (Fig. 2E-J).

### Weight error and stability

Fig. 2 quantifies how sample size affects CCA/PLS summary statistics. We next focused on error and stability of the weights, due to their centrality in CCA/PLS analyses in describing which features carry between-set association ^3^. Fig. 3A illustrates an example of how CCA/PLS weight vectors exhibit high error when typical sample-to-feature ratios are used. We systematically measured weight stability, i.e., the consistency of estimated weights across independent sample datasets, as a function of sample size. At small sample sizes, the average weight stability was close to 0 for CCA and eventually converged towards 1 (i. e. perfect similarity) with more samples (Fig. 3E). PLS exhibited striking differences from CCA: mean weight stability had a relatively high value even at low sample sizes, where weight error is very high (Figs. 3F, 2F) with high variability across population models.

**Figure 3.**
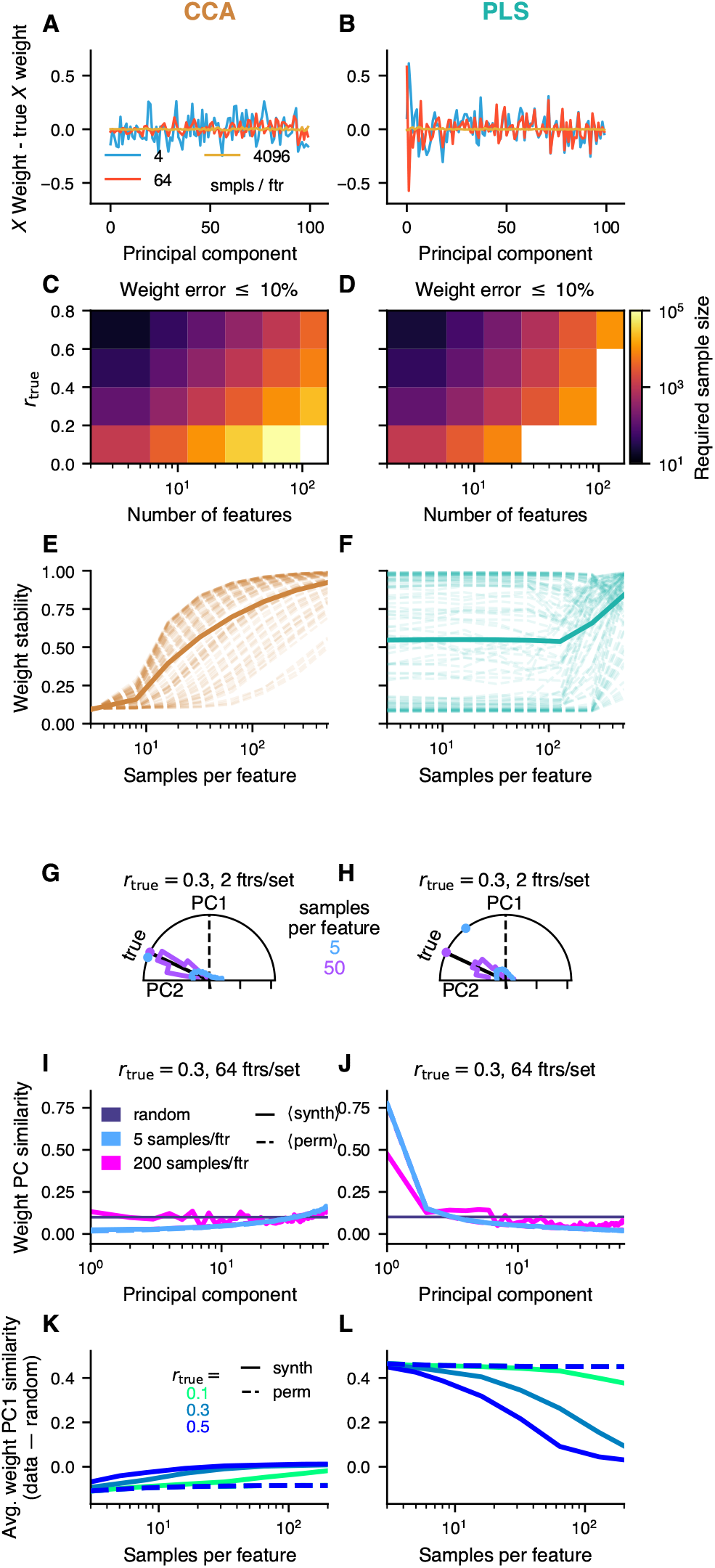
Large number of samples required to obtain good weight estimates. **A, B)** Realistic example where the true betwen-set correlation was set to *r*_true_ =0.3. Estimated weights are close to the assumed true (population) weights, as long as the sample size is large enough. **B)** For PLS even more samples were necessary. **C-D)** Sample sizes required to obtain less than 10% weight errors. **E-F)** Weight stability, i. e. the average cosine-similarity between weights across pairs of repetitions, increases towards 1 (identical weights) with more samples. For PLS, weight stability can be high, even with few samples. The true between-set correlation was set to *r*_true_=0.3. **G-H)** Example situation assuming a true between-set correlation of *r*_true_=0.3 between datasets and 2 features each for both *X* and *Y* datasets. In this 2-dimensional setting weight vectors, scaled to unit length, lie on a circle. Synthetic datasets were generated repeatedly. 5 samples per feature gave good estimates in many cases but notably all possible weight vectors occurred frequently. 50 sampes per feature resulted in consistently better estimates. Dots near border of semi-circles indicate directional means of distributions. **I-J)** Another example with 64 features per dataset and a between-set correlation *r*_true_=0.3. PLS weights have a strong PC1 similarity (cosine-similarity with first principal component). **K-L)** PC1 similarity was stronger for PLS (**L**) than for CCA (**K**) also for datasets with varying number of features and true between-set correlations *r*_true_. Shown is relative PC1 similarity across synthetic datasets with varying number of features, relative to the expected PC1 similarity of a randomly chosen vector with dimension matched to each synthetic dataset.

To assess the dependence of weight error on the assumed true between-set correlation and the number of features, we estimated the number of samples required to obtain less than 10% weight error (Fig. 3C-D). The required sample size is higher for increasing number of features, and lower for increasing true between-set correlation. We also observe that, by this metric, required sample sizes can be much larger than typical sample sizes in CCA/PLS studies.

### Weight PC1 similarity in PLS

At low sample sizes, PLS weights exhibit, on average, high error yet also relatively high stability (Figs. 3A-B,E-F and 2E-F). This suggests a systematic bias in PLS weights toward an axis different than the true latent axis of association. To gain further intuition, we first consider the illustrative case of each dataset comprising only 2 features, so that weight vectors are 2-dimensional unit vectors on a circle. With only 5 samples per feature (typical in CCA/PLS studies) (Fig. S3A), for CCA the circular histogram of weight vectors peaked around the true value, yet for PLS the peak was substantially shifted towards the first principal component axis (PC1).

We investigated how this weight bias toward PC1 in PLS manifests more generally. We quantified the PC similarity as the cosine similarity between estimated weight vectors and principal component axes, first with an illustrative data regime (64 features/dataset, *r*_true_ = 0.3). Compared to CCA, weight similarity to PC1 was strong for PLS, even with a large number of samples (Fig. 3I-J), and more so for a small number of samples. Finally, these observations also held for datasets from differing number of features and true correlations, with PLS weight vectors exhibiting strong bias toward PC1 especially for low sample sizes (Fig. 3L).

### Comparison of loadings and weights

In addition to weights, loadings provide a measure of importance for each considered variable ^32, 34^. We found that for CCA, stability and error of loadings followed a similar sample-size dependence as weights. In contrast, PLS loadings were extremely stable even at low sample sizes where error is high, indicating a strong bias (Fig. 4A,B). For both CCA and PLS, a steeper powerlaw describing the within-set variance produced more stable loadings (Fig. 4C,D). We next evaluated whether loadings exhibit bias toward principal component axes (Fig. 4E, F). At small sample sizes, PLS loadings and weights, as well as CCA loadings, strongly resembled more dominant principal component axes. Thus, the within-set variance can have strong biases on CCA/PLS results, irrespective of true between-set associations (Fig. S8).

**Figure 4.**
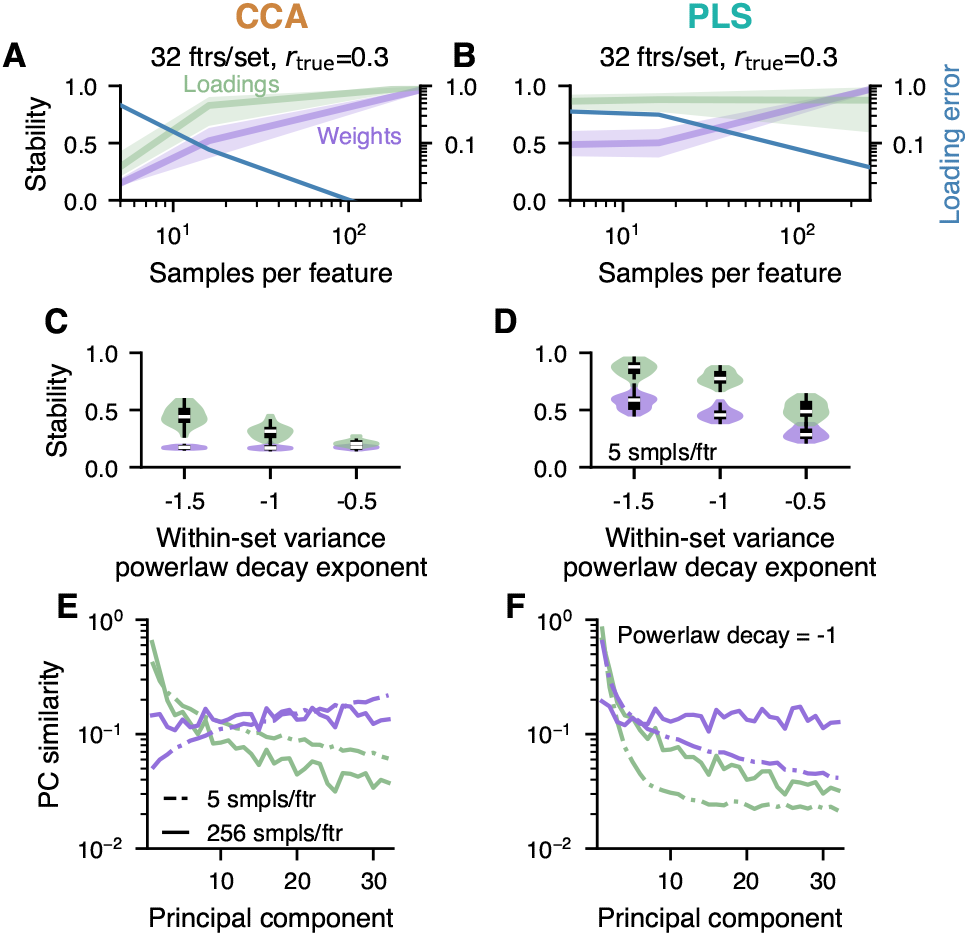
Stability and PC similarity of weights and loadings. Stability of loadings was quantified with pairwise Pearson correlation of loadings from independent samples, averaged over multiple generative models. **A,B)** PLS loadings exhibit very high stability despite large loading error at lower sample sizes, in contrast to CCA. **C,D)** The steeper the within-set variance spectrum the more stable were loadings for both CCA and PLS, and weights for PLS. **E,F)** CCA loadings resembled dominant principal component (PC) axes, while CCA weights for small sample sizes resembled more low-variance PC-axes. In contrast, PLS weights and loadings both exhibited strong bias toward dominant PC axes.

### Empirical brain-behavior CCA/PLS

Do these phenomena from our generative modeling framework hold in empirical data? We focused on two state-of-the-art population neuroimaging datasets: Human Connectome Project (HCP) ^1^ and UK Biobank (UKB) ^2^. Both provide multi-modal neuroimaging along with a wide range of behavioral and demographic measures, and both have been used for CCA-based brain-behavior mapping ^2, 3, 35–39^. HCP data is widely used and of cutting-edge quality, and the UKB is one of the largest publicly available population-level neuroimaging datasets.

We analyzed two modalities from the HCP, resting-state functional MRI (fMRI) (N=948) and diffusion MRI (dMRI) (N=1,020, Fig. S9A-D), and fMRI from the UKB (N=20,000). Functional and structural connectivity features were extracted from fMRI and dMRI, respectively. After modality-specific preprocessing (see Methods), datasets were deconfounded and reduced to 100 principal components (Fig. S1K), following prior CCA studies ^3, 35–39^. We repeatedly formed two non-overlapping subsamples of subjects, varying sizes up to 50 % of the subjects, and assessed CCA/PLS solutions (Figs. 5, S10, S11.).

**Figure 5.**
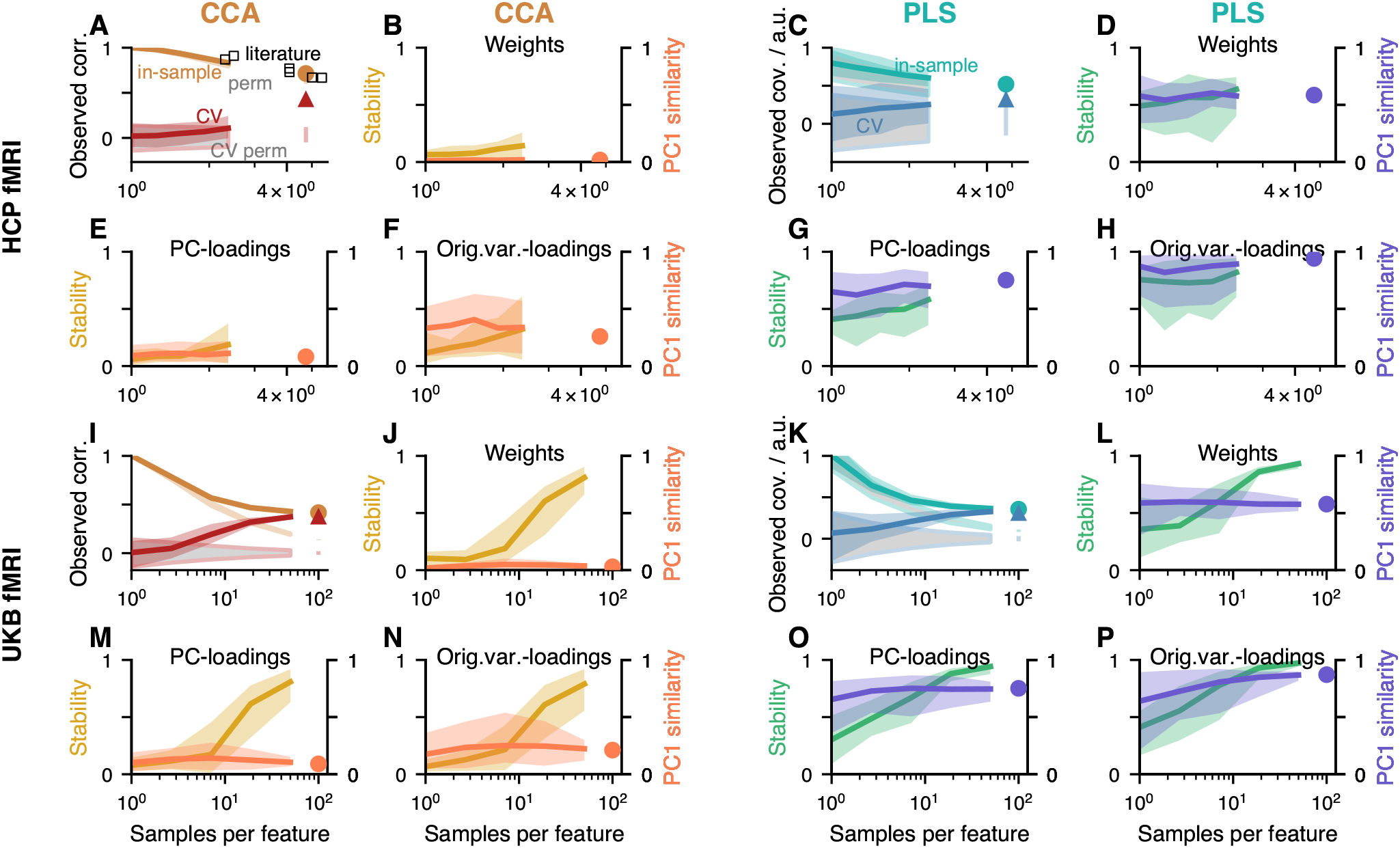
CCA and PLS analysis of empirical population neuroimaging datasets. For all datasets and for both CCA and PLS a significant mode of association was detected. Association strengths monotonically decreased with size of the subsamples (orange in column 1, green in column 3). Association strengths for permuted data are shown in grey (with orange and green outlines in columns 1 and 3, respectively). Deviations of the orange and green curves from the grey curves occur for sufficient sample sizes and correspond to significant *p*-values. Note how the curves clearly flatten for UKB but not for HCP data where the number of available subjects is much lower. Circle indicates the estimated value using all available data and the vertical bar in the same color below it denotes the corresponding 95 % confidence interval obtained from permuted data. In **A)** we also overlaid reported canonical correlations from other studies that used HCP data reduced to 100 principal components. Cross-validated association strengths shown in red (column 1) and blue (column 3), cross-validated estimation strengths of permuted datasets in grey with red and blue outlines in columns 1 and 3, respectively. Triangle indicates the cross-validated association strength using all data and the vertical bar in the same color below it denotes the corresponding 95 % confidence interval obtained from permuted data. Cross-validated association strengths were always lower than in-sample estimates and increased with sample size. For UKB (but not yet for HCP) cross-validated association strengths converged to the same value as the in-sample estimate. Weight stabilities increased with sample size for UKB and slightly for the PLS analyses of HCP datasets, while they remained low for the CCA analyses of HCP datasets. PC1 weight similarity was low for CCA but high for PLS. Both PC-loadings and original-variable-loadings show a similar pattern as weights, with loadings being slightly more similar to PC1 than weights. All analyses were performed with repeatedly subsampled data of varying sizes (*x*-axis). For each subsample size and repetition, we created two non-overlapping sets of subjects and calculated stability of weights / loadings using these non-overlapping pairs.

In-sample association strength decreased with increasing size of the subsamples, but converged to cross-validated association strength clearly only for the UKB at large sample size (Fig. 5A,C,I,K). Fig. 5A overlays reported CCA results from prior publications that used 100 features per set with HCP data, which further confirms the substantially decreasing association strengths as a function of sample size. HCP weight stabilities remained at low and intermediate values for CCA and PLS, respectively (Fig. 5B,D,J,L). In contrast, UKB weight stabilities reached values close to 1 (perfect stability). Moreover, for all datasets, PC1 similarity was close to 0 for CCA but markedly higher for PLS weights (Fig. 5B,D,J,L). Finally, loadings exhibited similar dependencies as weights, with higher PC1 similarity (Fig. 5E–H,M–P). Very similar behaviour is observed when using very different features extracted from diffusion MRI (Fig. S9).

All these empirical results are in agreement with analyses of synthetic data discussed above (Figs. 2-4). The overall similarity between CCA/PLS analyses of different neuroimaging modalities and features (Figs. 5, S9) suggests that sampling error is a major determinant in CCA/PLS solutions in typical data regimes. These results also show that stable CCA/PLS solutions with a large number of features can be obtained with UKB-size datasets.

We also explored reducing the data to different numbers of PCs than 100. Optimizing the number of PCs yielded 68 neural and 32 behavioral dimensions for HCP ^40^, which yielded higher cross-validated association strengths and higher stabilities of weights. In UKB, we separately varied the number of retained neuroimaging and behavioral principal components and calculated CCA/PLS association strengths (Fig. S12). We found that estimated association strengths rose strongly when retaining an increasing number of behavioral PCs, but only up to about 10. The situation for neuroimaging PCs differed between the methods, however. For CCA, retaining more neuroimaging PCs improved the association strength up to about 20-40 before plateauing. For PLS, on the other hand, the top PCs (*≈* 5-10) were enough for the association strength to plateau. Altogether, these results demonstrate the potential benefits of careful, modality-specific dimensionality reduction strategies to enhance CCA/PLS stability.

### Samples per feature alone predicts published CCA strengths

We next examined stability and association strengths in CCA analyses of empirical datasets more generally, through analysis of the published neuroimaging literature using CCA for brain-behavior associations. From 100 CCAs that were reported in 31 publications (see Methods), we extracted the number of samples, number of features, and association strengths. Most studies used less than 10 samples per feature (Fig. 6A and S3A). Overlaying reported canonical correlations as a function of samples per feature on top of predictions from our generative model shows that most published CCAs are compatible with a range of true correlations, from about 0.5 down to 0 (Fig. 6A). Remarkably, despite the variety of datasets and modalities used in these studies, the reported canonical correlation could be well predicted simply by the number of samples per feature alone (*R*^2^ = 0.83). We also note that reported CCAs might be biased upwards to some degree due to the fact that researchers might have explored a number of different analyses and reported the one with the highest canonical correlation.

**Figure 6.**
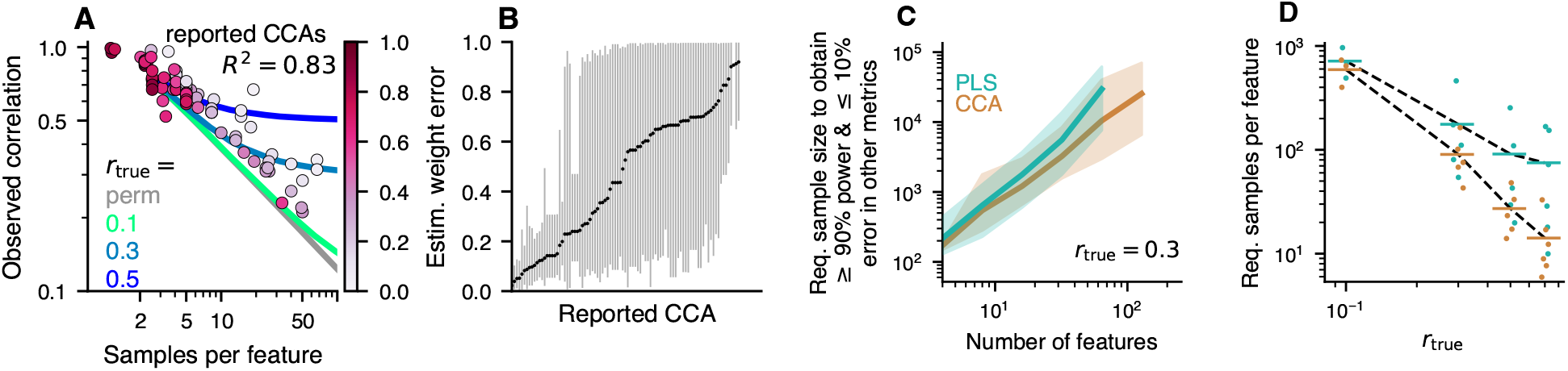
CCAs reported in the population neuroimaging literature might often be unstable. **A)** Canonical correlations and the number of samples per features are extracted from the literature and overlaid on predictions from the generative model for various between-set correlations *r*_true_. Many studies employed a small number of samples per feature (cf. also Fig. S3A) and reported a large canonical correlation. In fact, the reported canonical correlation can be predicted from the used number of samples per feature alone using linear regression (*R*^2^ = 0.83). We also estimated the weight error (encoded in the colorbar) for each reported CCA (details are illustrated in Fig. S13). **B)** The distribution of estimated weight errors for each reported CCA is shown along the *y*-axis. For many studies weight errors could be quite large, suggesting that conclusions drawn from interpreting weights might not be robust. **C-D)** Sample sizes to obtain at least 90 % power and at most 10 % error for the association strength, weight, scores and loadings. Shown estimates are constrained by the within-set variance spectrum (here *a_X_* + *a_Y_* = *−*2, cf. Fig. S16 for other values). **C)** Assuming a true between-set correlation of *r*_true_ = 0.3 (see Fig. S15A-D for other values) 100s to 1000s of samples are required to reach target power and error levels. Shaded areas show 95 % confidence intervals. **D)** The required number of samples divided by the total number of features in *X* and *Y* scales with *r*_true_. For *r*_true_ = 0.3 about 50 samples per feature are necessary to reach target power and error levels in CCA, which is much more than typically used (cf. Fig. S3A). Every point for a given *r*_true_ represents a different number of features and is slightly jittered for visibility. Values for a given dimensionality *p_X_* are only shown here if simulations were available for both CCA and PLS.

We next asked to what degree weight errors could be estimated from published CCAs. As these are unknown in principle, we estimated them using our generative modeling framework. We did this by (i) generating synthetic datasets of the same size as a given empirical dataset, and sweeping through assumed true correlations between 0 and 1, (ii) selecting those synthetic datasets for which the estimated canonical correlation matches the empirically observed one, and (iii) using the weight errors in these matched synthetic datasets as estimates for weight error in the empirical dataset (Fig. S13). This resulted in a distribution of weight errors across the matching synthetic datasets for each published CCA study that we considered. The mean of these distributions is shown in color overlay in Fig. 6A and the range of the distributions is shown in Fig. 6B (see also Fig. S3B). These analyses suggest that many published CCA studies likely have unstable feature weights due to an insufficient sample size.

### Calculator for required sample size

How many samples are required for stable CCA/PLS results, given particular dataset properties? One can base this decision on a combination of criteria, by bounding statistical power as well as relative error in association strength, weight error, score error and loading error at the same time. Requiring at least 90 % power and admitting at most 10 % error for other metrics, we determined the corresponding sample sizes in synthetic datasets by interpolating the curves in Fig. 2 (see Fig. S14A and Methods). The results are shown in Fig. 6C-D (see also Figs. S15, S16, S17). For example, when the true correlation is 0.3, several hundreds to thousands of samples are necessary to achieve the indicated power and error bounds (Fig. 6C). The required sample size per feature as a function of the true correlation roughly follows a powerlaw dependence, with a strong increase in required sample size when the true correlation is low (Fig. 6D). We also evaluated required sample sizes for a commonly used sparse CCA method (Fig. S18); however, an in-depth analysis of sparse CCA is beyond the scope of this study.

Finally, we formulated a concise, easy-to-use description of the relationship between model parameters and required sample size. To that end, we fitted a linear model to the logarithm of the required sample size, using logarithms of total number of features and true correlation as predictors (Fig. S14). We additionally included a predictor for the decay constant of the within-set variance spectrum, *|a_X_* + *a_Y_ |*. We found that a simple linear model approach yielded good predictive power for CCA and PLS, which we validated using split-half predictions (Fig. S14C-D).

## Discussion

Our generative modeling framework revealed how stability of CCA/PLS solutions depends on dataset properties. Our findings underscore that CCA/PLS stability and statistical significance do not need to coincide (see also ^41^). Importantly, estimated weight vectors, which govern the latent feature patterns that underlie an association, do not typically resemble the true weights when the number of samples is low (Fig. 3), which precludes generalizability and interpretability. PLS weights also show a consistent similarity to the first principal component axis (Fig. 3G-L), and therefore PLS weight stability is not sufficient to establish convergence to a true between-set relationship. The same pitfalls appear in state-of-the-art empirical datasets for brain-behavior associations.

CCA/PLS have become popular methods to reveal associations between neuroimaging and behavioral measures ^2, 3, 13, 20, 36–39, 42, 43^. The main interest lies in interpreting weights or loadings to understand the profiles of neural and behavioral features carrying the brain-behavior association. We have shown, however, that stability of weights and loadings are contingent on a sufficient sample size which, in turn, depends on the true between-set correlation. How strong are true between-set correlations for typical brain-behavior association studies? While this depends on the dataset at hand and is in principle unknown *a priori*, ^27^ report average cross-validated (out-of-sample) correlations of 0.17, whereas ^29^ argue that higher (*r >* 0.2) out-of-sample correlations are achievable with targeted methods. Our analyses provide insight to this question and highlight the importance of dataset dimensionality. We found in UKB convergence of canonical correlations to *∼*0.5. As the included behavioral measures comprised a wide assortment of categories, this canonical correlation is likely at upper end of expected ranges. Moreover, we found that most published brain-behavior CCA studies with substantially more than 10 samples per feature appeared to be compatible only with canonical correlations of *≤* 0.3, which is at the upper range suggested by recent empirical explorations ^27, 29^.

Assuming a relatively large between-set correlation of 0.3, our generative model still implies that *∼*50 samples per feature are required for stability of CCA solutions. For designs with hundreds of features, this necessitates many thousands of subjects, in agreement with ^27^. Many published brain-behavior CCAs do not meet this criterion. Moreover, in HCP data we saw clear signs that the available sample size was too small to obtain stable solutions—despite that the HCP is one of the largest and highest-quality neuroimaging datasets available to date. On the other hand, in the UKB, where we used 20000 subjects, CCA and PLS results converged with stability. As UKB-level sample sizes are well beyond what can be feasibly collected in typical neuroimaging studies, these findings support calls for aggregation of datasets that are shared widely ^44^.

For simplicity and tractability it was necessary to make a number of assumptions in our study. In particular, our synthetic data were normally distributed, which is typically not the case in practice. Importantly, empirical brain-behavior datasets yielded similar sample-size dependencies as synthetic datasets. We assumed the existence of a single cross-modality axis of association, whereas in practice several might be present.

Several related methods have been proposed to potentially circumvent shortcomings of standard CCA/PLS ^14^. Regularized or sparse CCA methods apply penalities to weight vectors to mitigate overfitting ^45^. We observed that its relative merit might depend on the true weight profile (S18). We note that a complete characterization of sparse CCA, and other methods such as non-linear extensions, was beyond the scope the present study. The numbers of features are important determinants for stability. Thus, methods for dimensionality reduction hold great promise. A variety of methods exist to determine an appropriate number of components for PCA and CCA ^40, 46–48^. Applying one method to HCP data yielded slightly better convergence (Fig. S9E-H). Alternatively, prior domain-specific knowledge could be used to preselect features hypothesized to be relevant for the question at hand.

In summary, we have presented a parameterized generative modeling framework for CCA and PLS. It allows analysis of the stability of CCA and PLS estimates, prospectively and retrospectively. We end by providing 9 recommendations for using CCA or PLS in practice (Table S1).

## Author contributions

Conceptualization: MH, SW, AA, SNS, JDM. Methodology: MH, JDM. Software: MH. Formal analysis: MH, SW, AM, BR. Resources: AA, SNS, JDM. Data Curation: AM, JLJ, AH. Writing - Original Draft: MH, JDM. Writing - Review & Editing: All authors. Visualization: MH. Supervision: JDM. Project administration: JDM. Funding acquisition: AA, SNS, JDM.

## Methods

### Experimental Design

The goal of this work was to determine requirements for stability of CCA and PLS solutions, both in simulated and empirical data. To do so, we first developed a generative model that allowed us to generate synthetic data with known CCA/PLS solutions. This made it possible to systematically study deviations of estimated from true solutions. Second, we used large state-of-the-art neuroimaging datasets with associated behavioral measurements to confirm the trends that we saw in synthetic data. Specifically, we used data from the Human Connectome Project (HCP) (n *≈* 1000) and UK Biobank (UKB) (n = 20000). Third, we analyzed published CCA results of brain-behavior relationships to investigate sample-size dependence of CCA results in the literature.

### Human Connectome Project (HCP) dataset

#### fMRI data

We used resting-state fMRI (rs-fMRI) from 951 subjects from the HCP 1200-subject data release (03/01/2017) ^1^. The rs-fMRI data were preprocessed in accordance with the HCP Minimal Preprocessing Pipeline (MPP). The details of the HCP preprocessing can be found elsewhere ^50, 51^. Following the HCP MPP, BOLD time-series were denoised using ICA-FIX ^52, 53^ and registered across subjects using surface-based multimodal inter-subject registration (MSMAll) ^54^. Additionally, global signal, ventricle signal, white matter signal, and subject motion and their first-order temporal derivatives were regressed out ^55^.

The rs-fMRI time-series of each subject comprised of 2 (69 subjects), 3 (12 subjects), or 4 (870 subjects) sessions. Each rest session was recorded for 15 minutes with a repetition time (TR) of 0.72 s. We removed the first 100 time points from each of the BOLD sessions to mitigate any baseline offsets or signal intensity variation. We subtracted the mean from each session and then concatenated all rest sessions for each subject into a single time-series. Voxel-wise time series were parcellated to obtain region-wise time series using the “RelatedValidation210” atlas from the S1200 release of the HCP ^56^. Functional connectivity was then computed as the Fisher-*z*-transformed Pearson correlation between all pairs of parcels. 3 subjects were excluded (see below), resulting in a total of 948 subjects with 64620 connectivity features each.

#### dMRI data

Diffusion MRI (dMRI) data and structural connectivity patterns were obtained as described in ^57, 58^. In brief, 41 major white matter (WM) bundles were reconstructed from preprocessed HCP diffusion MRI data ^59^ using FSL’s XTRACT toolbox ^58^. The resultant tracts were vectorised and concatenated, giving a WM voxels by tracts matrix. Further, a structural connectivity matrix was computed using FSL’s probtrackx ^60, 61^, by seeding cortex/white-grey matter boundary (WGB) vertices and counting visitations to the whole white matter, resulting in a WGB *×* WM matrix. Connectivity “blueprints” were then obtained by multiplying the latter with the former matrix. This matrix was parcellated (along rows) into 68 regions with the Desikan-Killany atlas ^62^ giving a final set of 68 *×* 41 = 2788 connectivity features for each of the 1020 HCP subjects.

#### Behavioral measures

The same list of 158 behavioral and demographic data items as in ^3^ was used.

#### Confounds

We used the following items as confounds: Weight, Height, BPSystolic, BPDiastolic, HbA1C, the third cube of FS BrainSeg Vol, the third cube of FS IntraCanial Vol, the average of the absolute as well as the relative value of the root mean square of the head motion, squares of all of the above, and an indicator variable for whether an earlier of later software version was used for MRI preprocessing. Head motion and software version were only included in the analysis of fMRI vs behavioral data, not in the analysis of dMRI vs behavioral data. Confounds were inverse-normal-transformed (ignoring missing values) such that each had mean 0. Subsequently, missing values were set to 0. 3% and 5% of confound values were missing in the fMRI vs. behavior, and dMRI vs behavior analysis, respectively. All resulting confounds were *z*-scored once more.

### UK Biobank (UKB) dataset

#### fMRI data

We utilized pre-processed resting-state fMRI data ^64^ from 20,000 subjects, available from the UK Biobank Imaging study ^2^.

In brief, EPI unwarping, distortion and motion correction, intensity normalization and high-pass temporal filtering were applied to each subject’s functional data using FSL’s Melodic ^66^, data were registered to standard space (MNI), and structured artefacts are removed using ICA and FSL’s FIX ^52, 53, 66^. A set of resting-state networks were identified, common across the cohort using a subset of subjects (*≈* 4000 subjects) ^64^. This was achieved by extracting the top 1200 components from a group-PCA ^67^ and a subsequent spatial ICA with 100 resting-state networks ^66, 68^. Visual inspection revealed 55 non-artifactual ICA components. Next, these 55 group-ICA networks were dual regressed onto each subject’s data to derive representative timeseries for each of the ICA components. Following the regression of the artifactual nodes for all other nodes and the subsequent removal of the artifactual nodes, the timeseries were used to compute partial correlation parcellated connectomes with a dimensionality of 55 *×* 55. The connectomes were z-score transformed and the upper triangle vectorized to give 1485 functional connectivity features per subject, for each of the 20,000 subjects.

#### Behavioral measures

The UK Biobank contains a wide range of subject measures ^69^, including physical measures (e.g., weight, height), food and drink, cognitive phenotypes, lifestyle, early life factors and sociodemographics. We hand-selected a subset of 389 cognitive, lifestyle and physical measures, as well as early life factors. For categorical items, we replaced negative values with 0, as in ^2^. Such negative values encode mostly “Do not know”/“Prefer not to answer”. Measures with multiple visits were then averaged across visits, reducing the number of measures 226. We then performed a check for measures that had missing values in more than 50% of subjects and also for measures that had identical values in at least 90% of subjects; no measures were removed through these checks. We then performed a redundancy check.

Specifically, if the correlation between any two measures was *>* 0.98, one of the two items was randomly chosen and dropped. This procedure further removed 2 measures, resulting in a final set of 224 behavioral measures, available for each of the 20000 subjects.

#### Confounds

We used the following items as confounds: acquisition protocol phase (due to slight changes in acquisition protocols over time), scaling of T1 image to MNI atlas, brain volume normalized for head size (sum of grey matter and white matter), fMRI head motion, fMRI signal-to-noise ratio, age and sex. In addition, similarly to ^2^, we used the squares of all non-categorical items (i. e. T1 to MNI scaling, brain volume, fMRI head motion, fMRI signal-to-noise ratio and age), as well as age *×* sex and age^2^*×* sex.

Altogether these were 14 confounds. Confounds were inverse-normal-transformed (ignoring missing values), such that each had mean 0. 6% of values were missing and set to 0. All resulting confounds were then *z*-scored across subjects once more.

### Preprocessing of empirical data for CCA and PLS

We prepared data for CCA following, for the most part, the pipeline in Ref. 3.

#### Deconfounding

Deconfounding of a matrix *X* with a matrix of confounds *C* was performed by subtracting linear predictions, i.e.

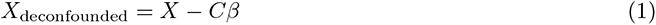

where

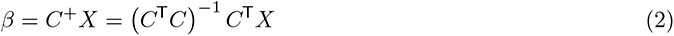

The confounds used were specific to each dataset and mentioned in the previous section.

#### Neuroimaging data

Neuroimaging measures were *z*-scored. The resulting data matrix was de-confounded (as described above), decomposed into principle components via a singular value decomposition, and the left singular vectors, multiplied by their respective singular values, were used as data matrix *X* in the subsequent CCA or PLS analysis. We retained 100 principal components (out of 948, 1020 and 1485 for the HCP-fMRI, HCP-dMRI and UKB analysis, respectively).

#### Behavioral and demographic data

The list of used behavioral items were specific to each dataset and mentioned in the previous sections. Given this list, separately for each item, a rank-based inverse normal transformation ^70^ was applied and the result *z*-scored. For both of these steps subjects with missing values were disregarded. Next, a subjects *×* subjects covariance matrix across variables was computed, considering for each pair of subjects only those variables that were present for both subjects. The nearest positive definite matrix of this covariance matrix was computed using the function `cov_nearest` from the Python statsmodels package ^71^. This procedure has the advantage that subjects can be used without the need to impute missing values. An eigenvalue decomposition of the resulting covariance matrix was performed where the eigenvectors, scaled to have standard deviation 1, are principal component scores. They are then scaled by the square-roots of their respective eigenvalues (so that their variances correspond to the eigenvalues) and used as matrix *Y* in the subsequent CCA or PLS analysis. We retained 100 (out of 948, 1020 and 20000 for the HCP-fMRI, HCP-dMRI and UKB analysis, respectively) principal components corresponding to the highest eigenvalues.

### Generating synthetic data for CCA and PLS

Note that mathematical derivations and additional explanation of terminology related to CCA and PLS are provided in the Supporting Information document. We analyzed properties of CCA and PLS with simulated datasets from a multivariate generative model. These datasets are drawn from a normal distribution with mean 0 and a covariance matrix Σ that encodes assumed relationships in the data. To specify Σ we need to specify relationships of features within *X*, i. e. the covariance matrix Σ*_XX_ ∈* R*^pX ×pX^*, relationships of features within *Y*, i. e. the covariance matrix Σ*_Y Y_ ∈* R*^pY ×pY^*, and relationships between features in *X* on the one side and *Y* on the other side, i.e. the matrix Σ*_XY_ ∈* R*^pX ×pY^*. Together, these three covariance matrices form the joint covariance matrix (Fig. 1D)

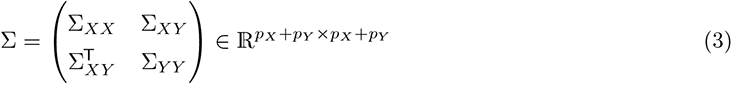

for *X* and *Y* and this allows us to generate synthetic datasets by sampling from the associated normal distribution *N* (0, Σ). *p_X_* and *p_Y_* correspond to the number of features in *X* and *Y* respectively.

### The covariance matrices Σ*_XX_* and Σ*_Y Y_*

Given a data matrix *X*, the features can be re-expressed in a different coordinate system through multiplication by an orthogonal matrix *O*: *X̃* = *XO*. No information is lost in this process, as it can be reversed: *X* = *X̃O*^T^. Therefore, we are free to make a convenient choice. We select the principal component coordinate system as in this case the covariance matrix becomes diagonal, i. e. Σ*_XX_* = diag(*σ⃗_XX_*). Analogously, for *Y* we choose the principal component coordinate system such that Σ*_Y Y_* = diag(*σ⃗_Y Y_*). For modeling, to obtain a concise description of *σ⃗_XX_* and *σ⃗_Y Y_* we assume a power-law such that *σ_XX,i_* = *c_XX_i^−aXX^* and *σ_Y Y,i_* = *c_Y Y_ i^−aY Y^* with decay constants *a_XX_* and *a_Y Y_* (Fig. 1B). Unless a match to a specific dataset is sought, the scaling factors *c_XX_* and *c_Y Y_* can be set to 1 as they would only rescale all results without affecting conclusions.

### The cross-covariance matrix **Σ*_XY_***

#### PLS

Given a cross-covariance matrix Σ*_XY_*, PLS solutions can be derived via a singular value decomposition as

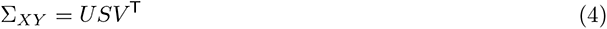

where the singular vectors in the columns of *U* and *V* are orthonormal and the matrix *S* contains the singular values on its diagonal. The columns of *U* and *V* are the weight vectors for *X* and *Y*, respectively, and the singular values give the corresponding between-set association strengths. Thus, the above equation allows us to compute Σ*_XY_* given orthonormal PLS weight vectors *U*, *V* and corresponding between-set association strengths diag(*S*). This is what we do for PLS. See below for how we select the specific weights that we use. Singular values here in the context of PLS are covariances between *X* scores and *Y* scores. We re-express these in terms of a correlation, i.e. the *i*-th singular value, *s_i_*, is

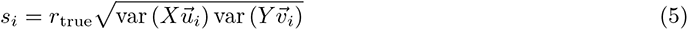

where *r*_true_ is the assumed true (population) canonical correlation for mode *i*, and *u⃗_i_* and *v⃗_i_*are the *i*-th columns of *U* and *V*, respectively.

#### CCA

For CCA we know that a “whitened” version of the between-set covariance matrix is related to weight vectors and canonical correlations via the singular value decomposition as

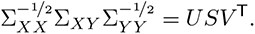

As above, the singular vectors in the columns of *U* and *V* are orthonormal and the matrix *S* contains the singular values on its diagonal. For CCA, the singular values directly give the canonical correlations, but the singular vectors are not identical to the weight vectors. Instead, CCA weights, *W_X_* and *W_Y_* for *X* and *Y*, repectively, are given by

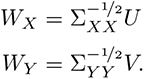

Thus, given weight vectors as columns of *W_X_* and *W_Y_*, as well as population canonical correlations diag(*S*), we can calculate the corresponding between-set covariance matrix for CCA as

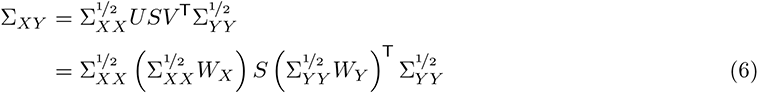

Importantly, we had assumed that the columns of *U*, as well as *V*, were orthonormal. Thus, the columns of *W_X_* and *W_Y_* must satisfy the following constraints: 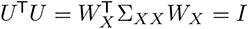, as well as 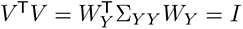. One straightforward way to do so, is to restrict our consideration to one mode, i. e. we assume that Σ*_XY_* has rank one (or can be approximated with a rank one matrix) such that *U*, *V*, as well as *W_X_* and *W_Y_* consist of only 1 column. In this case, whatever weight vector we choose, say 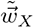 (analogously for *Y*), we can normalize it as

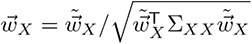

and *w⃗_X_* will satisfy the constraint. See below, for how we select the specific weight vectors we use in our simulations.

### Choice of weight vectors

As weight vectors, we choose random unit vectors of the desired dimension, as long as they satisfy the following two constraints:

1. We aim to obtain association modes that explain a “large” amount of variance in the data, otherwise the resulting scores could be strongly affected by noise. The decision is based on the explained variance of only the first mode and we require that it is greater than ^1^*/*2 of the average explained variance of a principal component in the dataset, i.e. we require that

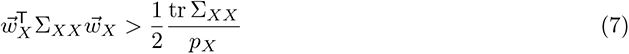

and analogously for *Y*.

2. The weight vectors impact the joint covariance matrix Σ (via (3), (4) and (6)). Therefore, we require that the chosen weights result in a proper, i. e. positive definite, covariance matrix Σ.

### Summary

Thus, to generate simulated data for CCA and PLS, we vary the assumed between-set correlation strengths *ρ--_XY_*, setting them to select levels, while choosing random weights *W_X_* and *W_Y_*. The columns of the weight matrices *W_X_* and *W_Y_* must be mutually orthonormal for PLS, while for CCA they must satisfy 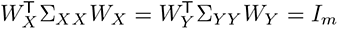.

### Performed simulations

For Figs. 2, 3C-D, the colored curves in Fig. 6A, Figs. S7E-F, 6C-D, S15A-D, and the left 3 columns of Fig. S2, we ran simulations for *m* = 1 between-set association mode assuming true correlations of 0.1, 0.3, 0.5, and 0.7, used dimensionalities *p_X_* = *p_Y_* of 2, 4, 8, 16, 32, and 64 as well as 25 different covariance matrices (10 for *p_X_* = *p_Y_* = 64 in combination with *r*_true_ = 0.7). *a_X_* + *a_Y_* was fixed at -2. 100 synthetic datasets were drawn from each instantiated normal distribution. Where not specified otherwise, null distributions were computed with 1000 permutations. Due to computational expense, some simulations did not finish and are reported as blank spaces in heatmaps.

Similar parameters were used for other figures, except for the following deviations.

For Fig. 3A-B *p_X_*was 100, *r*_true_ = 0.3, *a_X_* = *a_Y_* = *−*1 and we used 1 covariance matrix for CCA and PLS.

For Fig. 3E-F *p_X_* was 100, *r*_true_ = 0.3 and we used 100 different covariance matrices.

For Fig. 3G-H, *p_X_* was 2, *r*_true_ = 0.3, *a_X_* = *a_Y_* = *−*1 and we used 10000 different covariance matrices for CCA and PLS.

For Fig. 3I-L, we used 2, 4, 8, 16, 32 and 64 for *p_X_*, 0.1, 0.3 and 0.5 for *r*_true_, 10 different covariance matrices for CCA and PLS, and 10 permutations. A subset of these, namely *p_X_* = 64 and *r*_true_ = 0.3 was used for Fig. 3I-J.

For Fig. 4, *p_X_* was 32, *r*_true_ =0.3, *a_X_* = *a_Y_* was -1.5, -1.0 and -0.5, and we used 25 different covariance matrices.

For Fig. 6, we varied *r*_true_ from 0 to 0.99 in steps of 0.01 for each combination of *p_X_* and *p_Y_* for which we have a study in our database of reported CCAs, assumed *a_X_* = *a_Y_* = 0, and generated 1 covariance matrix for each *r*_true_.

For the right 3 columns in Fig. S2 *p_X_* + *p_Y_* was fixed at 64, for *p_X_* we used 2, 4, 8, 16, 32 and 40 different covariance matrices.

In Fig. S5, for *p_X_* we used 4, 8, 16, 32, 64, we generated 10 different covariance matrices for both CCA and PLS and varied *r*_true_ from 0 to 0.99 in steps 0.01.

For Fig. S6 we used 2, 4, 8, 16 and 32 for *p_X_*, and 10 different covariance matrices for both CCA and PLS.

For Fig. S16E-L, Fig. S14, and Fig. S17 we used 25 different covariance matrices. For each instantiated joint covariance matrix, *a_X_* + *a_Y_* was chosen uniformly at random between -3 and 0 and *a_X_* was set to a random fraction of the sum, drawn uniformly between 0 and 1.

In Fig. S18 we used *p_X_* = *p_Y_* =64, *r*_true_ =0.3, *a_X_* = *a_Y_* =-1.0 and 4 different covariance matrices.

### Meta-analysis of prior literature

A PubMed search was conducted on December 23, 2019 using the query (`“Journal Article”[Publication Type]) AND (fmri[MeSH Terms] AND brain[MeSH Terms]) AND (“canonical correlation analysis”`) with filters requiring full text availability and studies in humans. In addition, studies known to the authors were considered. CCA results were included in the meta-analysis if they related neuroimaging derived measures (e. g. structural or functional MRI, . . .) to behavioral or demographic measures (e. g. questionnaires, clinical assessments, . . .) across subjects, if they reported the number of subjects and the number of features of the data entering the CCA analysis, and if they reported the observed canonical correlation. This resulted in 100 CCA analyses reported in 31 publications, which are summarized in SI Dataset 1.

### The *gemmr* software package

We provide an open-source Python package, called *gemmr*, that implements the generative modeling framework presented in this paper https://github.com/murraylab/gemmr. Among other functionality, it provides estimators for CCA, PLS and sparse CCA; it can generate synthetic datasets for use with CCA and PLS using the algorithm laid out above; it provides convenience functions to perform sweeps of the parameters on which the generative model depends; and it calculates required sample sizes to bound power and other error metrics as described above. For a full description, we refer to the package’s documentation.

## Statistical Analysis

### Evaluation of sampling error

We use five metrics to evaluate the effects of sampling error on CCA and PLS analyses.

#### Statistical power

Power measures the capability to detect an existing association. It is calculated when the true correlation is greater than 0 as the probability across 100 repeated draws of synthetic datasets from the same normal distribution that the observed association strength (i. e. correlation for CCA, covariance for PLS) of a dataset is statistically significant. Significance is declared if the *p*-value is below *α* = 0.05. The *p*-value is evaluated as the probability that association strengths are greater in the null-distribution of association strengths. The corresponding null-distribution is obtained from performing CCA or PLS on 1000 datasets where the rows of *Y* were permuted randomly. Power is bounded between 0 and 1 and, unlike for the other metrics (see below), higher values are better.

#### Relative error in between-set covariance

The relative error of the between-set association strength is calculated as

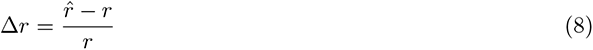

where *r* is the true between-set association strength and *r*^ is its estimate in a given dataset.

#### Weight error

Weight error Δ*w* is calculated as 1 - absolute value of cosine similarity between observed 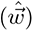 and true (*w--*) weights, separately for data sets *X* and *Y*, and the greater of the two errors is taken:

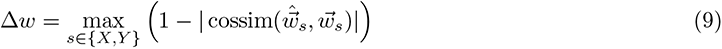

where

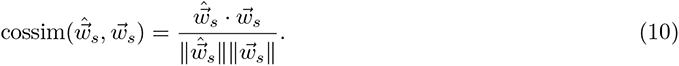

The absolute value of the cosine-similarity is used due to the sign ambiguity of CCA and PLS. This error metric is bounded between 0 and 1 and measures the cosine of the angle between the two unit vectors 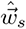 and *w⃗_s_*.

#### Score error

Score error Δ*t* is calculated as 1 – absolute value of Pearson correlation between observed and true scores. The absolute value of the correlation is used due to the sign ambiguity of CCA and PLS. As for weights, the maximum over datasets *X* and *Y* is selected:

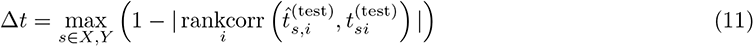

Each element of the score vector represents a sample (subject). Thus, to be able to compute the correlation between estimated 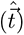 and true (*t⃗*) score vectors, corresponding elements must represent the same sample, despite the fact that in each repetition new data matrices are drawn in which the samples have completely different identities. To overcome this problem and to obtain scores, which are comparable across repetitions (denoted 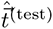 and *t⃗*^(test)^), each time a set of data matrices is drawn from a given distribution *N* (0, Σ) and a CCA or PLS model is estimated, the resulting model (i. e. the resulting weight vectors) is also applied to a “test” set of data matrices, *X*^(test)^ and *Y* ^(test)^ (of the same size as *X* and *Y*) obtained from *N* (0, Σ) and common across repeated dataset draws. The score error metric Δ*t* is bounded between 0 and 1.

#### Loading error

Loading error Δ*ℓ* is calculated as (1 *−* absolute value of Pearson correlation) between observed (i.e., sample) and true (i.e., population) loadings. The absolute value of the correlation is used due to the sign ambiguity of CCA and PLS. As for weights, the maximum over datasets *X* and *Y* is selected:

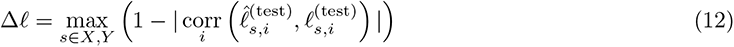

True loadings are calculated with SI Eq. 3 (replacing the sample covariance matrix in the formula with its population value). Estimated loadings are obtained by correlating data matrices with score vectors (SI Eq. 2). Thus, the same problem as for scores occurs: the elements of estimated and true loadings must represent the same sample. Therefore, we calculate loading errors with loadings obtained from test data (*X*^(test)^ and *Y* ^(test)^) and test scores (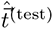 and *t⃗*^(test)^) that were also used to calculate score errors.

The loading error metric Δ*ℓ* is bounded between 0 and 1 and reflects the idea that loadings measure the contribution of original data variables to the between-set association mode uncovered by CCA and PLS.

Loadings are calculated by correlating scores with data matrices. All synthetic data matrices in this study are based in the principal component coordinate system. In practice, however, this is not generally the case. Nonetheless, as the transformation between principal component and original coordinate system cannot be constrained, here we do not consider this effect.

### Weight similarity to principal component axes

The directional means *µ* in Fig. 4A-B are obtained via

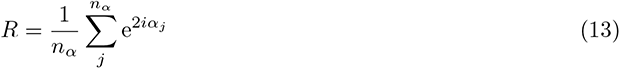

as *µ* = arg(*R*)*/*2.

To interpret the distribution of cosine similarities between weights and the first principal component axis we compare this distribution to a reference, namely to the distribution of cosine-similarities between a random *n*-dimensional unit vector and an arbitrary other unit vector *e⃗*. This distribution *f* is given by:

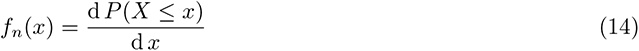

where *P* denotes the cumulative distribution function for the probability that a random unit-vector has cosine-similarity with *e⃗* (or, equivalently, projection onto *e⃗*) *≤ x*. For *−*1 *≤ x ≤* 0, *P* can be expressed in terms of the surface area *A_n_*(*h*) of the *n*-dimensional hyperspherical cap of radius 1 and height *h* (i. e.

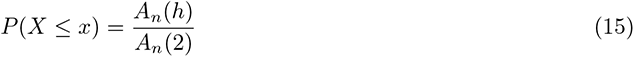

where *A_n_*(2) is the complete surface area of the hypersphere and

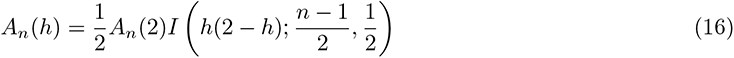

and *I* is the regularized incomplete beta function. Thus,

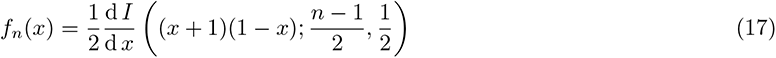

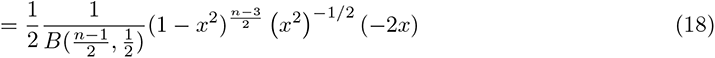

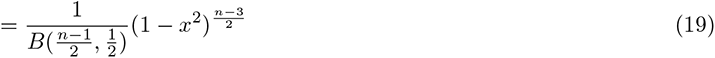

where *B* is a beta function and

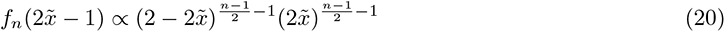

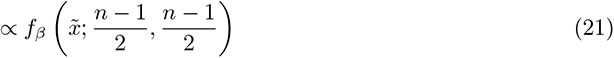

where *f_β_* is the probability density function for the beta distribution. Hence, 2*X̃ −* 1 with 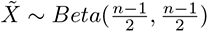 is a random variable representing the cosine similarity between 2 random vectors (or, equivalently, the projection of a random unit-vector onto another).

### CCA/PLS analysis of empirical data

Permutation-based *p*-values in Fig. 5 and S9 were calculated as the probability that the CCA or PLS association strength of permuted datasets was at least as high as in the original, unpermuted data.

Specifically, to obtain the *p*-value, rows of the behavioral data matrix were permuted and each resulting permuted data matrix together with the unpermuted neuroimaging data matrix were subjected to the same analysis as the original, unpermuted data, in order to obtain a null-distribution of between-set associations. 1000 permutations were used.

Due to familial relationships between HCP subjects they are not exchangeable so that not all possible permutations of subjects are appropriate ^72^. To account for that, in the analysis of HCP fMRI vs behavioral data, we have calculated the permutation-based *p*-value as well as the confidence interval for the whole-data (but not the subsampled data) analysis using only permutations that respect familial relationships. Allowed permutations were calculated using the functions hpc2blocks and palm quickperms with default options as described in https://fsl.fmrib.ox.ac.uk/fsl/fslwiki/PALM/ExchangeabilityBlocks (accessed May 18, 2020). No permutation indices were returned for 3 subjects that were therefore excluded from the functional connectivity vs behavior analysis.

Subsampled analyses (Fig. 5) were performed for 5 logarithmically spaced subsample-sizes between 202 and 50% of the total subject number. For each subsample size 100 pairs of non-overlapping data matrices were used.

Cross-validated analyses were performed with 5-fold cross-validation.

### Principal component spectrum decay constants

The decay constant of a principal component spectrum (Fig. S1A-J) was estimated as the slope of a linear regression (including an intercept term) of log(explained variance of a principal component) on log(principal component number). For each dataset in Fig. S1A-J we included as many principal components into the linear regression as necessary to explain either 30% or 90% of the variance.

### Determination of required sample size

As all evaluation metrics change approximately monotonically with the samples per feature ratio, we fit splines of degree 3 to interpolate and to determine the number of samples per feature that approximately results in a given target level for the evaluation metric. For power (higher values are better) we target 0.9, for all the other metrics (lower values are better) we target 0.1. Before fitting the splines, all samples per feature are log-transformed and metrics are averaged across repeated datasets from the same covariance matrix. Sometimes the evaluation metrics show non-monotonic behavior and in case the cubic spline results in multiple roots we filter those for which the spline fluctuates strongly in the vicinity of the root (suggesting noise), and select the smallest remaining root *ñ* for which the interpolated metric remains within the allowed error margin for all simulated *n > ñ*, or discard the synthetic dataset if all roots are filtered out. In case a metric falls within the allowed error margin for all simulated *n* (i. e. even the smallest simulated *n*_0_) we pick *n*_0_.

We suggest, in particular, a *combined* criterion to determine an appropriate sample size. This is obtained by first calculating samples per feature sizes with the interpolation procedure just described separately for the metrics power, relative error of association strength, weight error, score error and loading error. Then, for each parameter set, the maximum is taken across these five metrics.

### Sample-size calculator for CCA and PLS

Estimating an appropriate sample size via the approach described in the previous section is computationally expensive as multiple potentially large datasets have to be generated and analyzed. To abbreviate this process (see also Fig. S14A) we use the approach from the previous section to obtain sample-size estimates for *r*_true_ *∈ {*0.1, 0.3, 0.5, 0.7, 0.9}, *p_X_* ∈ {2, 4, 8, 16, 32, 64, 128}, *p_Y_* = *p_X_*, and *a_X_* + *a_Y_ ∼ U* (*−*3, 0), *a_X_* = *c*(*a_X_* + *a_Y_*), and *c ∼ U* (0, 1), where *U* denotes a uniform distribution. We then fit a linear model to the logarithms of the sample size, with predictors log(*r*_true_), log(*p_X_* + *p_Y_*), *|a_X_* + *a_Y_ |*, and including an intercept term.

We tested the predictions of linear model using a split-half approach (Fig. S14B-F), i. e. we refitted the model using either only sample-size estimates for *r*_true_ *∈ {*0.1, 0.3} and half the values for *r*_true_ = 0.5, or the other half of the data, and tested the resulting refitted model on the remaining data in each case.

### Data availability

Human Connectome Project and UK Biobank datasets cannot be made publicly available due to data use agreements. Human Connectome Project and UK Biobank are available for researchers to apply for data access. The outcomes of synthetic datasets that were analyzed with CCA or PLS are available from https://osf.io/8expj/.

### Code availability

Our open-source Python software package, gemmr, will be freely available at https://github.com/murraylab/gemmr. Jupyter notebooks detailing the analyses and generation of figures presented in the manuscript will be made available as part of the package documentation.

---

**Table S1.** Considerations and recommendations for using CCA and PLS in practice.

| # | Keyword | Recommendation |
| --- | --- | --- |
| 1. | Importance of sample size and number of features | Sample size and the number of features in the dataset are of critical importance for the stability of CCA and PLS. Dimensionality reduction (e.g. PCA) is a useful preprocessing step, as long as it does not remove components correlated between sets. Methods for selecting number of components that take into account the correlation between sets have been proposed, e.g.<sup>40</sup>, or with the difficulties of interpreting CCA results in mind<sup>47</sup>. |
| 2. | Significance testing | A significant non-zero association does not necessarily indicate that estimated weights are reliable. |
| 3. | Association strength error | In-sample estimates for association strengths are too high, cross-validated estimates too low, their average tended to be better. |
| 4. | Weights & loadings | Weights and loadings estimated with too few samples are unreliable. For PLS, estimation of cross-loadings required fewer samples than loadings. |
| 5. | PC1 similarity | In PLS, weights can appear consistently similar to the first principal component axis. |
| 6. | Deceptive weight stability | For PLS, weights can appear stable, scattering around the first principal component axis, and converge to their true values only for very large sample sizes. |
| 7. | Subsampling | Subsampling can be used to check stability of estimated association strengths in empirical data: similar results for varying subsample sizes indicate stability. |
| 8. | Reporting | Number of samples, number of features (after dimensionality reduction) and obtained association strength (in-sample and cross-validated) should be reported. The within-set variance spectrum is useful as well. |
| 9. | Required sample size | Generally, we recommend at least 50 samples per feature for CCA, more for PLS (depending on the variance spectrum). The accompanying Python package (*GEMMR*) can be used to calculate recommended sample sizes for given dataset characteristics. |

**Figure S1.**
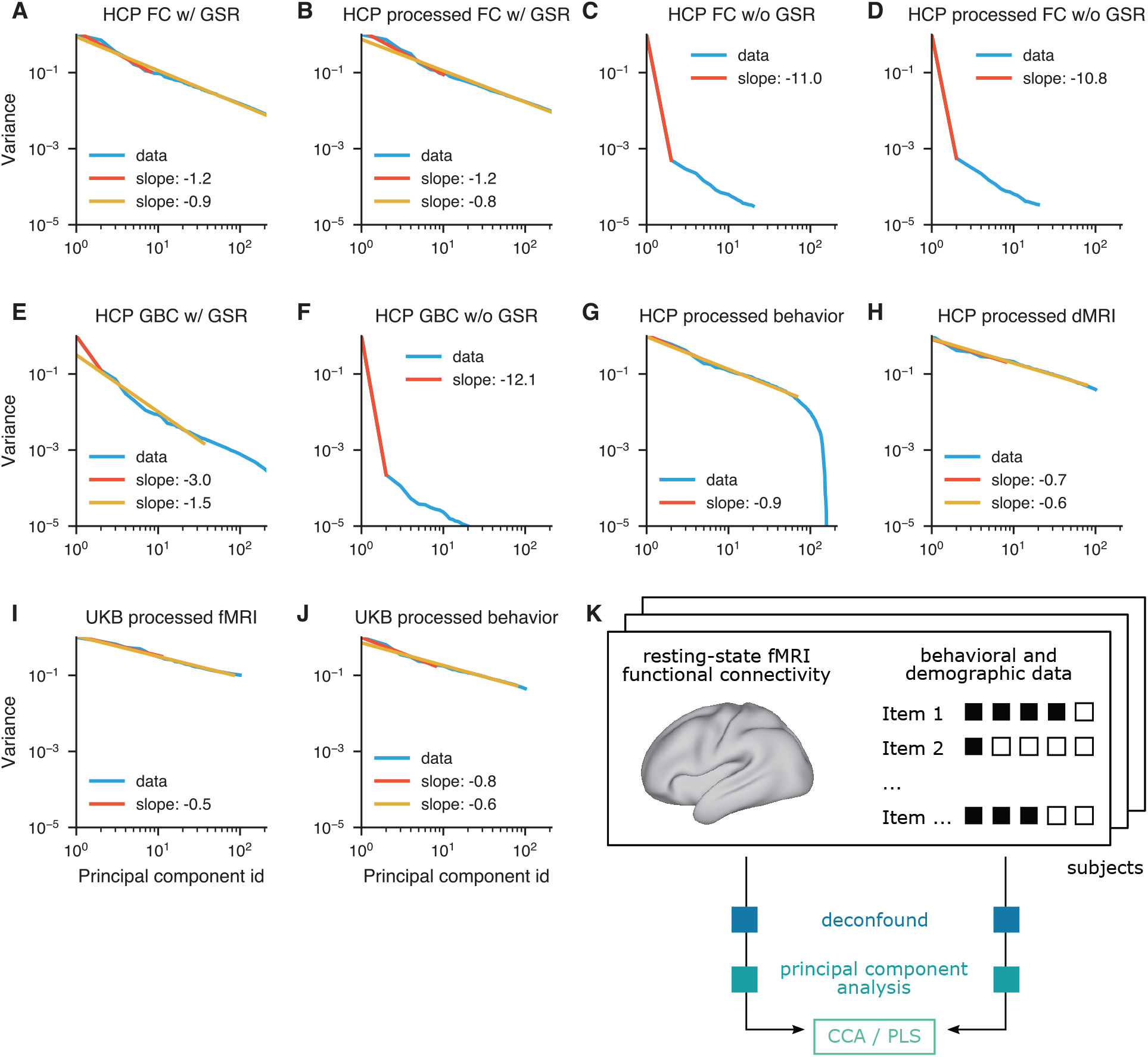
Supplementary analyses of empirical data. **A-J)** Decay con_C_s_C_t_A_an_/_ t_P_s_LS_of principal component spectra in empirical data. Decay constants are estimated as the slope in a linear regression for the logarithm of the explained variance on the logarithm of the associated principal component number. We include enough components into the linear regression as necessary to explain either 30 % (red) or 90 % (yellow) of the variance. Where the two resulting slopes coincide only one is shown. Shown are decay constants for the following data matrices: **A)** HCP functional connectivity and **B)** HCP functional connectivity after preprocessing for CCA / PLS (as described in subsection), both based on 951 subjects. **C)** HCP functional connectivity for 877 subjects where global signal was not regressed out (cf. subsection) and **D)** HCP functional connectivity of 877 subjects where global signal was not regressed out after preprocessing for CCA / PLS. **E)** HCP global brain connectivity (GBC), i. e. the sum across rows of the parcel *×* parcel functional connectivity matrix (951 subjects) and **F)** HCP GBC where global signal was not regressed out (877 subjects). **G)** HCP behavioral data of 951 subjects after preprocessing for CCA / PLS **H)** HCP diffusion MRI structural connectivity blueprints of 1020 subjects after preprocessing for CCA / PLS. **I)** UK Biobank fMRI of 20000 subjects after preprocessing for CCA / PLS, **J)** UK Biobank behavioral measures of 20000 subjects after preprocessing for CCA / PLS. **K)** HCP data analysis workflow. Resting-state functional connectivity data and behavioral and demographic data from corresponding subjects were separately deconfounded, reduced to 100 principal components and then analyzed with CCA and PLS.

**Figure S2.**
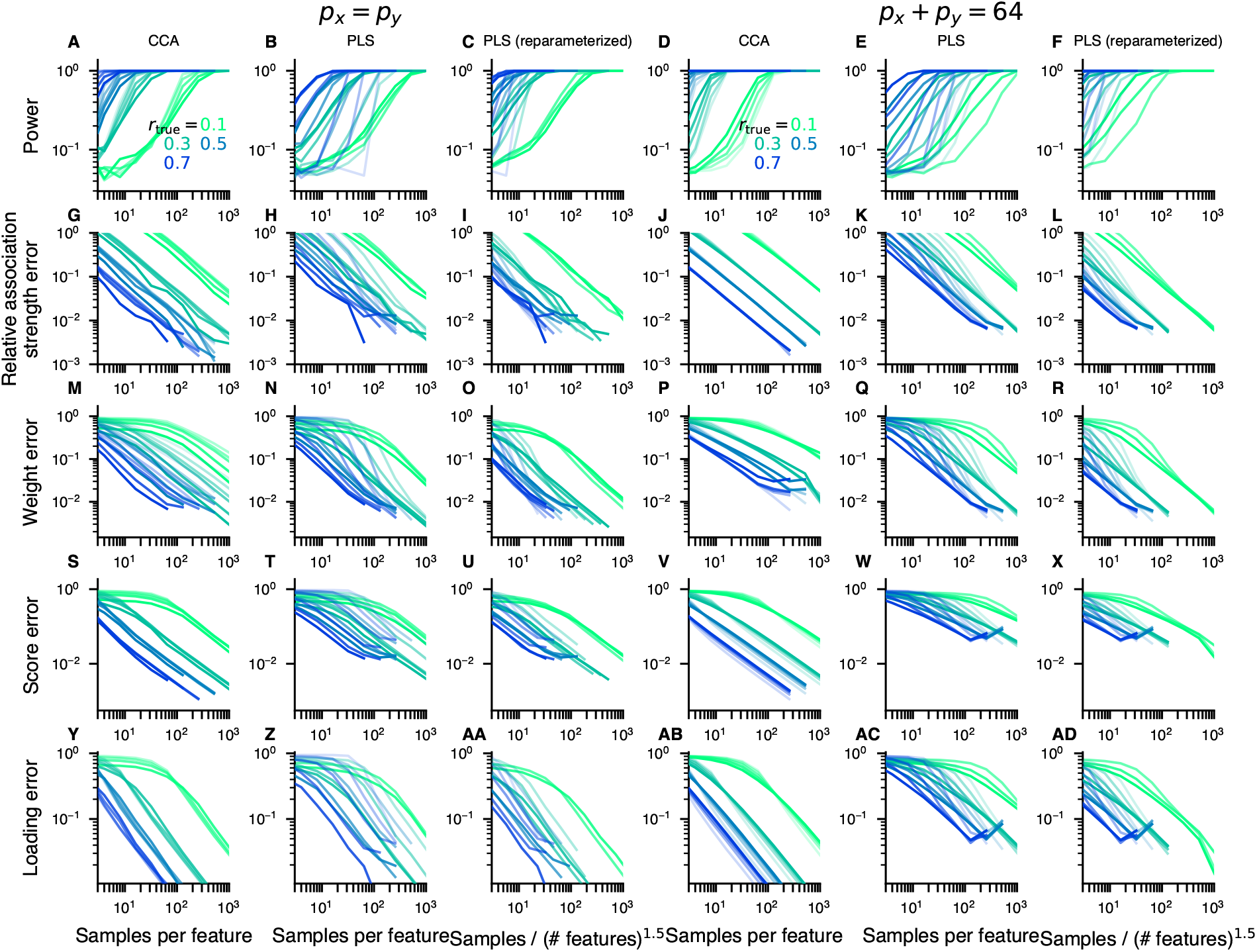
Samples per feature is a key effective parameter. Throughout the manuscript we have presented results in terms of the parameter “samples per feature”. Here, we demonstrate that this is, approximately, a good parameterization. *p_X_* and *p_Y_* denote the number of features for *X* and *Y* respectively. Color hue represents true between-set correlation *r*_true_, saturated colors are used for *p_X_* = 2, and fainter colors for higher *p_X_* (in this figure, *p_X_* ∈ {2, 4, 8, 16, 32, 64}), where *p_X_* (*p_Y_*) refers to the number of features in the *X* (*Y*) dataset. We fixed *p_X_* = *p_Y_* in the left 3 columns, whereas we fixed *p_X_* + *p_Y_* = 64 (and thus had *p_X_* ≠ *p_Y_*) in the right 3 columns. In CCA (first column), for a given *r*_true_, power and error metric curves for various number of features are very similar when parameterized as “samples per feature”. In PLS (second column), the same tendency can be observed, albeit the overlap between curves of the same hue (i. e. with same *r*_true_ but different number of features) is worse. When “samples / (number of features)^1.5^” is used instead (third column), the curves overlap more. The same trends can be seen in the right 3 columns, where *p_X_* ≠ *p_Y_*.

**Figure S3.**
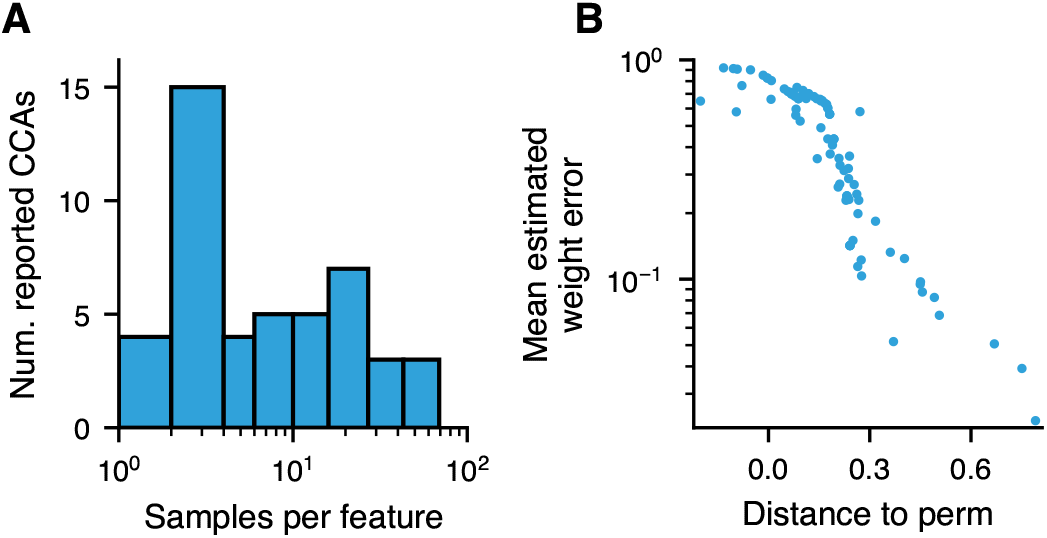
Supplementary results related to analysis of prior literature. **A)** Typical number of samples per feature in brain-behavior CCAs. Studies using CCA to analyze brain-behavior relationships often used less than 5 samples per feature. Note that we here considered the number of features that entered into the CCA analysis, which, after preprocessing, can be considerably less than the “raw” number of features. **B)** Distance from null in *samples-per-feature vs observed correlation* plot predicts weight error. A linear model was fit to the simulated, permuted data shown in Fig. 6A and for each reported CCA the orthogonal distance to the fit-line was measured and is shown here on the *x*-axis, with positive values indicating deviations towards the top-right corner of Fig. 6A. The mean estimated weight error for the reported CCAs is smaller the farther away from the permuted data the CCA lies in the top-right part of the plot.

**Figure S4.**
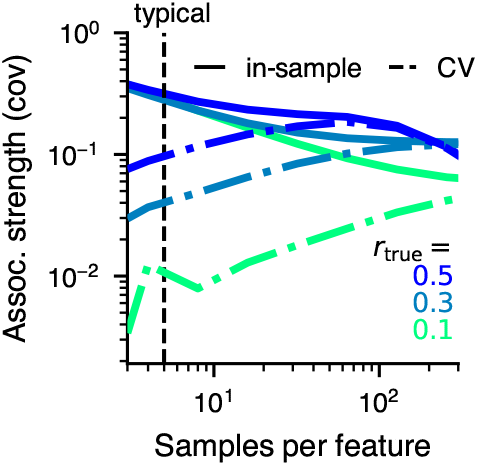
Sample-size dependence of between-set covariance for PLS. In-sample estimates (solid lines) decrease, while cross-validated estimates increase with samples-per-feature.

**Figure S5.**
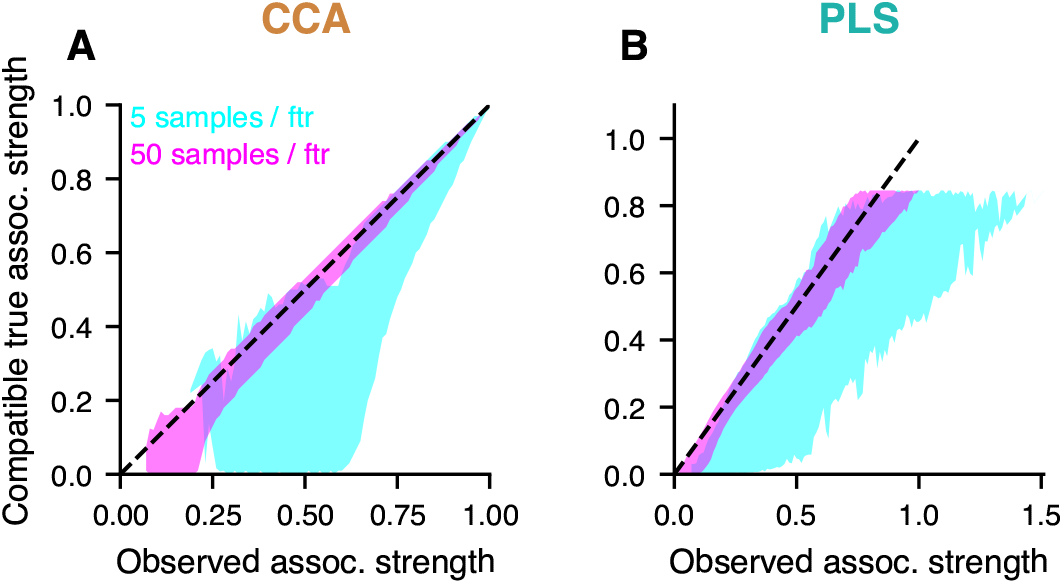
A wide range of true association strengths is compatible with a given observed association strength. Synthetic datasets were generated where the true correlation was varied from 0 to 0.99 in steps of 0.01 and analyzed with **A)** CCA, **B)** PLS. We investigated 4, 8, 16, 32, 64 and 128 features per set, set up 10 different covariance matrices with differing true weight vectors for each number of features and true correlation, and drew 100 repeated datasets from each corresponding normal distribution. For every CCA and PLS we recorded the observed association and binned them in bins with width 0.01. The plots show 95 % confidence intervals of the true association strength that were associated with a given observed association strength. Notably, apart from the very strongest observed association strengths which indicate an almost equally strong true correlation, compatible true association strengths can be markedly lower, down to essentially 0, when the number of used samples per feature is low.

**Figure S6.**
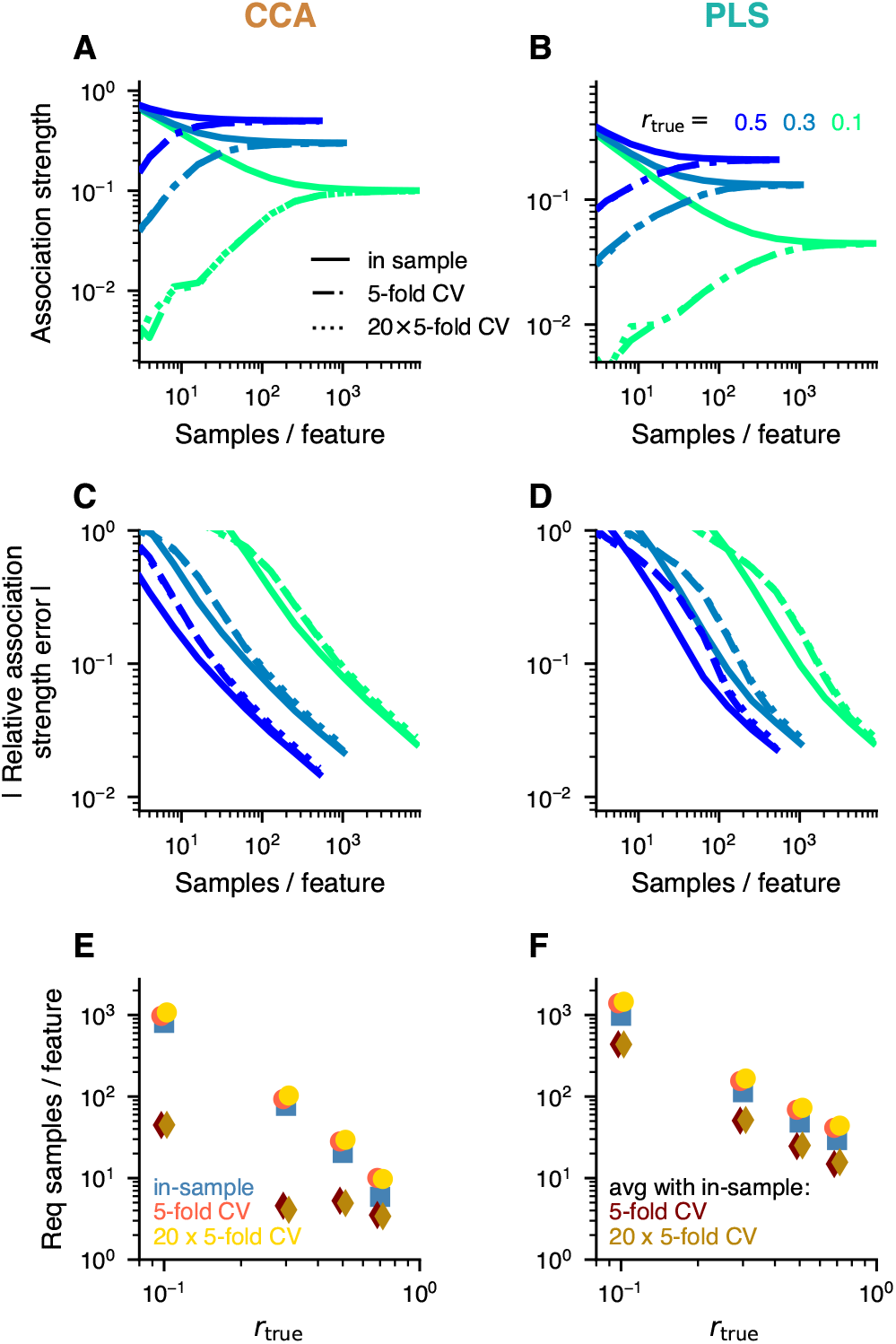
Cross-validated estimation of association strength. In contrast to in-sample estimates, cross-validated estimates of between-set association strengths underestimate the true value *r*_true_. We tested two different cross-validation strategies here with very similar results (curves overlap): 5-fold cross-validation (dash-dotted line) and a strategy where the data were randomly split 20 times into 80 % train and 20 % test (“20*×*5-fold CV”, dotted line). **C-D)** The absolute value of the relative estimation error is similar for in-sample and cross-validated estimates. **E-F)** Using the average of the in-sample and cross-validated estimates results in a better estimate than either of those, so that less samples are required to reach a target error level (here: 10 %).

**Figure S7.**
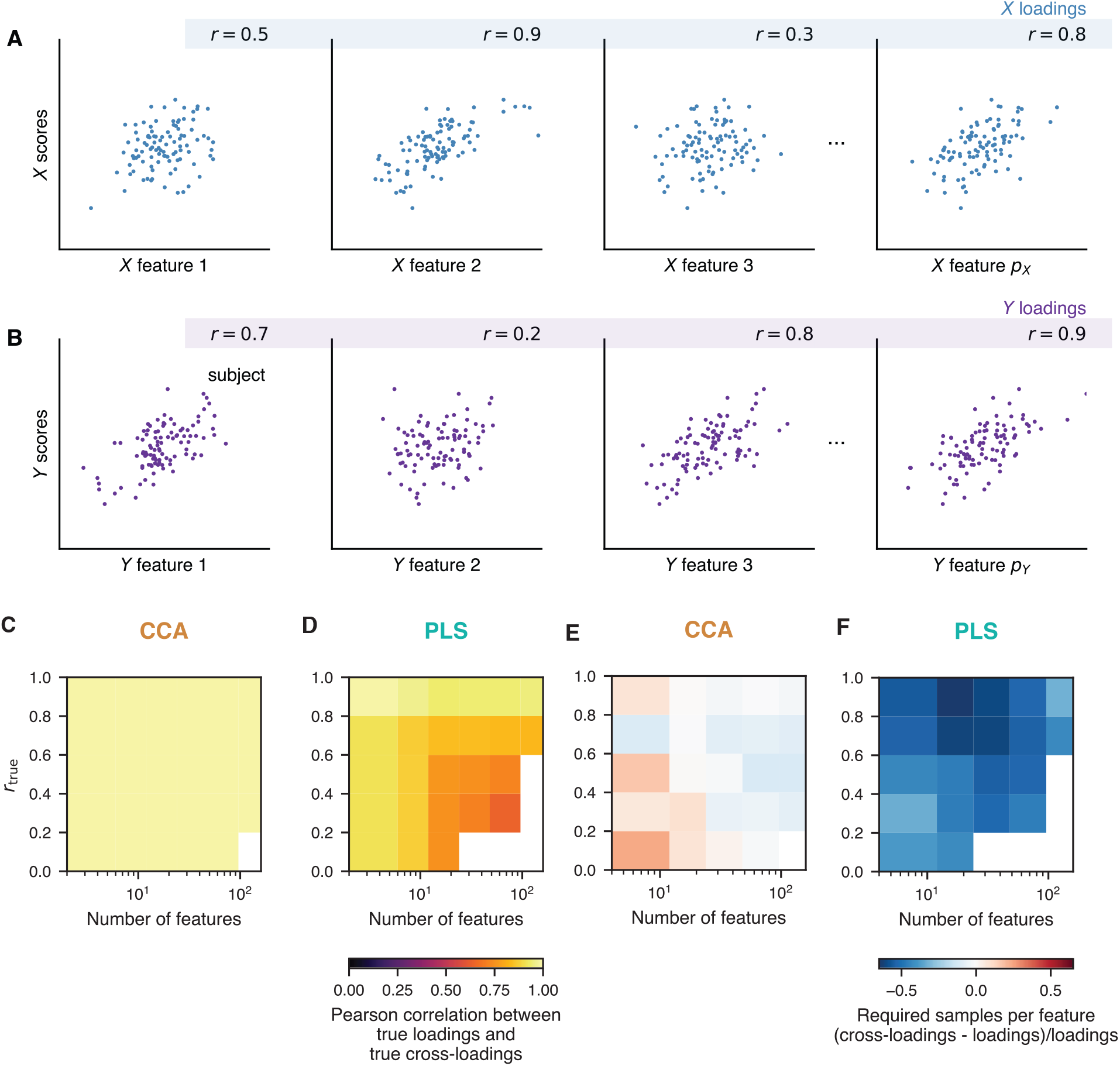
Loadings and cross-loadings. **A-B)** Loadings are defined as Pearson correlations across samples of a feature with the CCA/PLS scores. The loadings vector contains these correlations for all variables. Apart from the illustrated loadings, *cross-loadings* in which scores of one set are correlated with the original features of the other set can also be computed. **C-D)** True loadings and cross-loadings. **C)** In CCA, true loadings and true cross-loadings were collinear. **D)** For PLS, they were strongly correlated. The shown correlations were averaged across 25 covariance matrices with different true weight vectors. *a_X_* + *a_Y_* was constrained to -2. **E-F)** For PLS cross-loadings provide more stable estimates of feature profiles than loadings. Samples-per-feature required to obtain less than 10% error in either loadings or cross-loadings are compared. Shown here is their relative difference, i. e. the required sample-per-features for cross-loadings minus for loadings, divided by the required samples-per-feature for loadings. **E)** Relative differences were small for CCA. **F)** However, for PLS less samples were required with cross-loadings than with loadings to obtain the same error level. *r*_true_ indicates the true between-set correlations used in each respective simulation.

**Figure S8.**
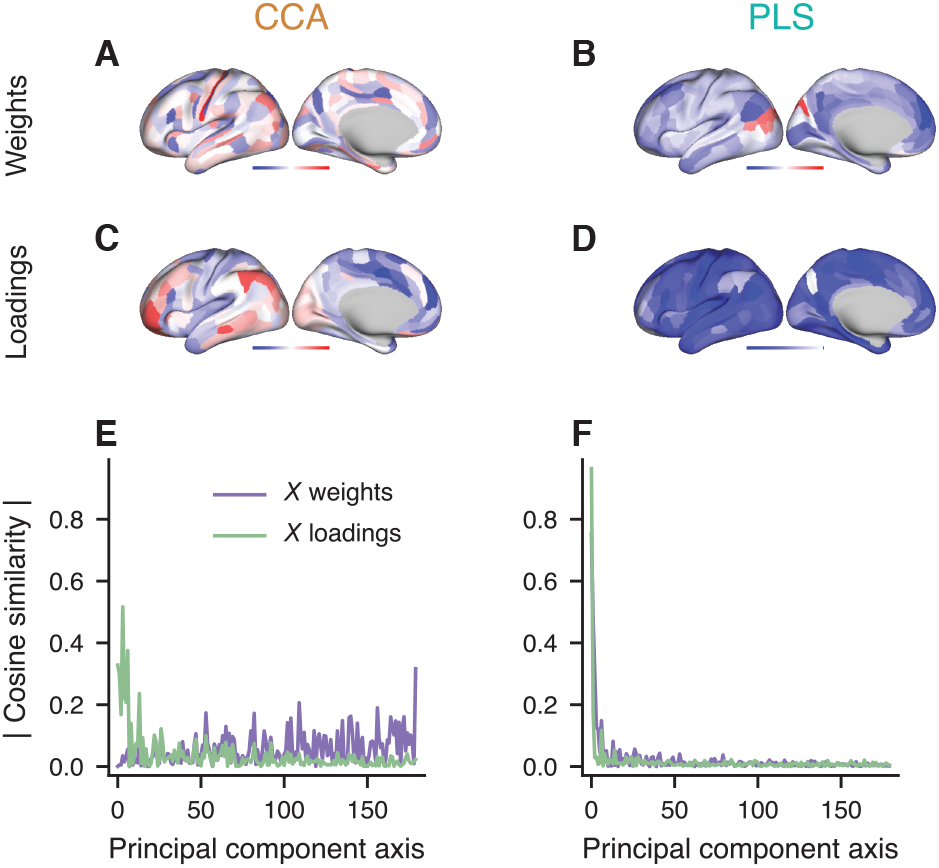
Weights vs loadings in real data. Using 180 fMRI-GBC features and the 5 dominant behavioral principal components as input to CCA / PLS we here illustrate GBC weights and loadings. **A** CCA weights, **B** PLS weights, **C** CCA loadings, and **D** PLS loadings. Note the relative noisiness of CCA weights. **E-F** shows a decomposition of weights and loadings into principal components, illustrating that CCA weights overlap more with low-variance PC-axes, while CCA loadings, as well as PLS weights and loadings overlap more with dominant PC-axes.

**Figure S9.**
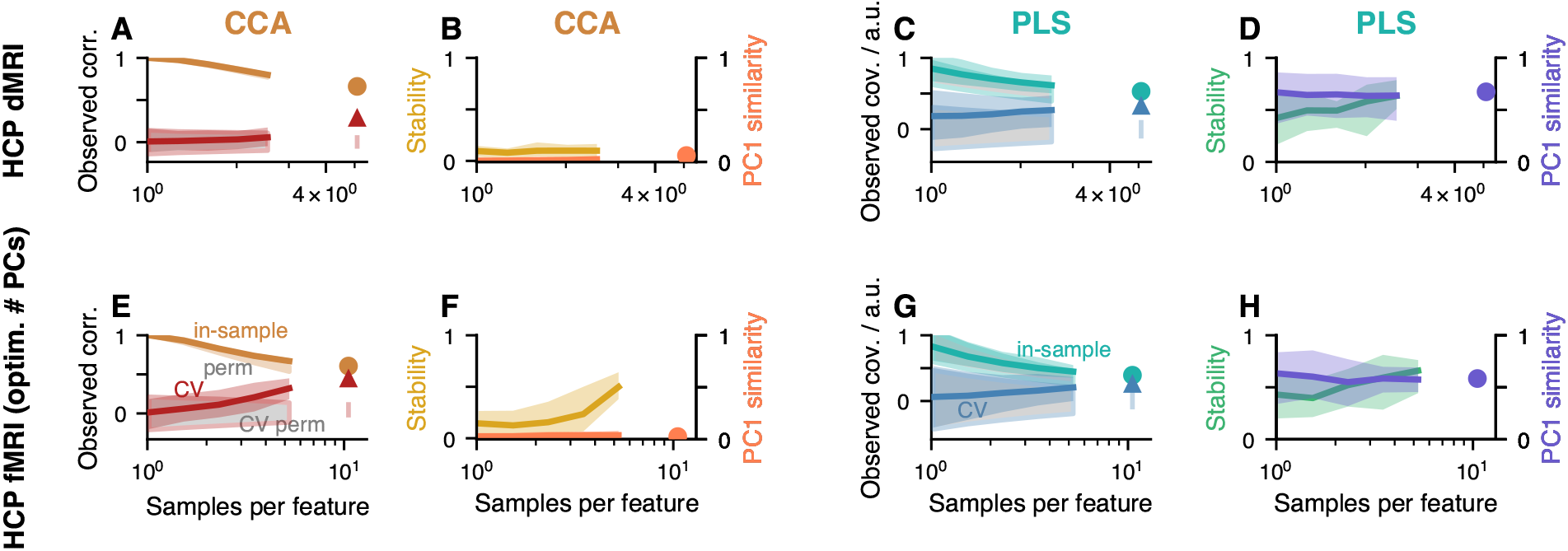
Additional CCA and PLS analyses of HCP data. Layout is similar to first row in Fig. 5. **A-D**) HCP dMRI data was related to behavioral and demographic data. Overall, CCA and PLS behave similarly using dMRI compared to fMRI data (Fig. 5A-D). *p*-values in **A** and **C** were 0.001 and 0.001, respectively. **E-H)** Re-analysis of HCP fMRI vs behavior data with optimized number of principal components. Format is identical to Fig. 5. The only difference is the number of principal components retained for analysis: whereas in Fig. 5 100 principal components were used for both datasets, in agreement with previous studies of HCP data, here we chose the number of principal component with a “max-min detector”. As the algorithm provided multiple values for the optimal number of components *p_X_* (neuroimaging data) and *p_Y_* (behavioral and demographic data), we selected here the pair that minimized *p_X_* + *p_Y_*. The optimized values were *p_X_* = 59 and *p_Y_* = 31, along with 13 between-set modes (we only consider the first one here). *p*-values for CCA and PLS were, respectively, 0.001 and 0.004. While the results are very similar to Fig. 5, (i) the observed correlations in **E)** appear to have stabilized more and are lower than in Fig. 5A, (ii) in-sample and cross-validated association strengths are more similar here in panels **A)** and **C)** than in Fig. 5, and (iii) weight similarities in **B)** and **D)** are higher than in Fig. 5. Altogether results seem to have converged more with the same sample size. This demonstrates the potential benefit of dimensionality reduction for CCA and PLS.

**Figure S10.**
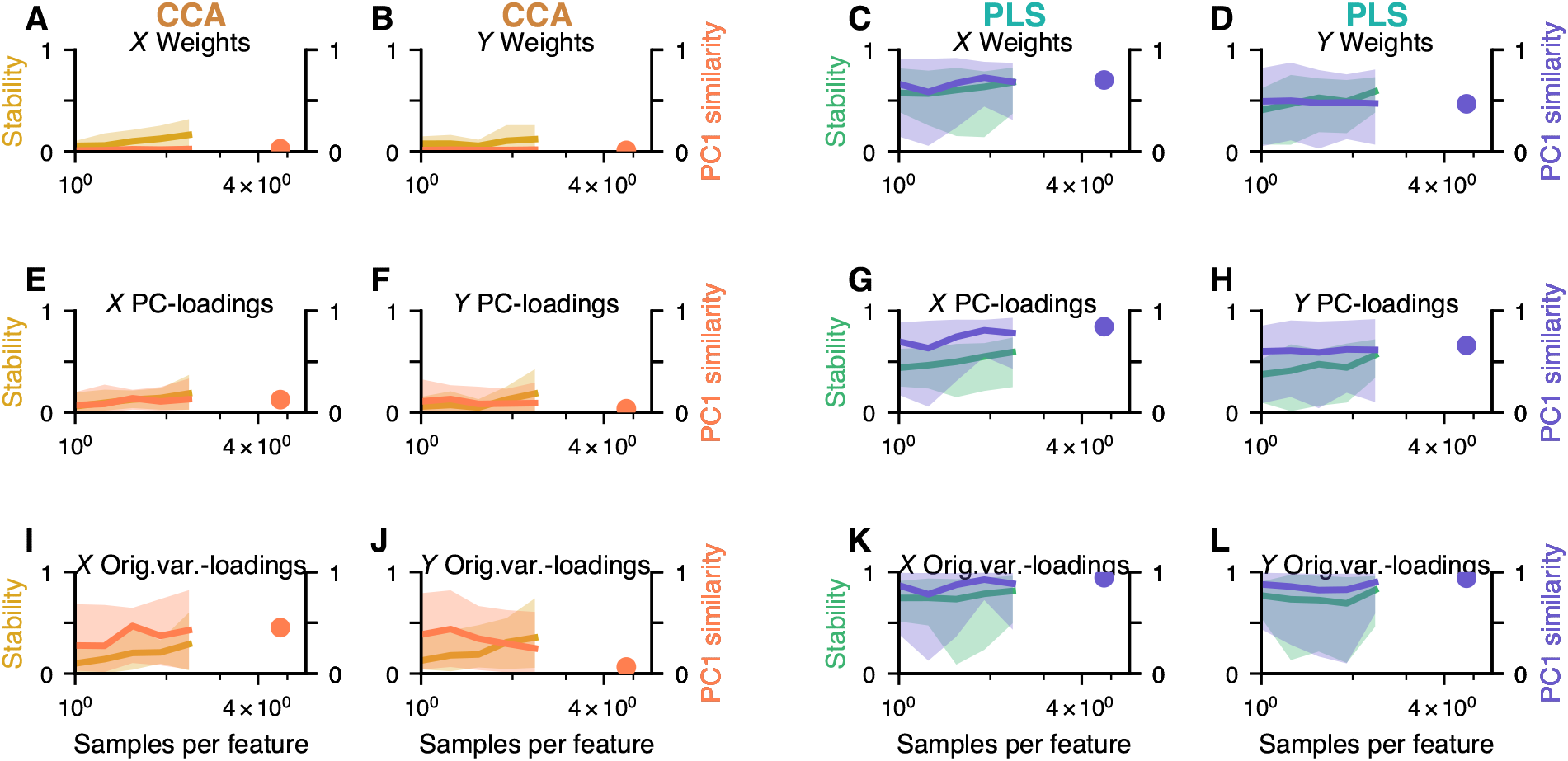
HCP fMRI CCA/PLS results for *X* and *Y* separately.

**Figure S11.**
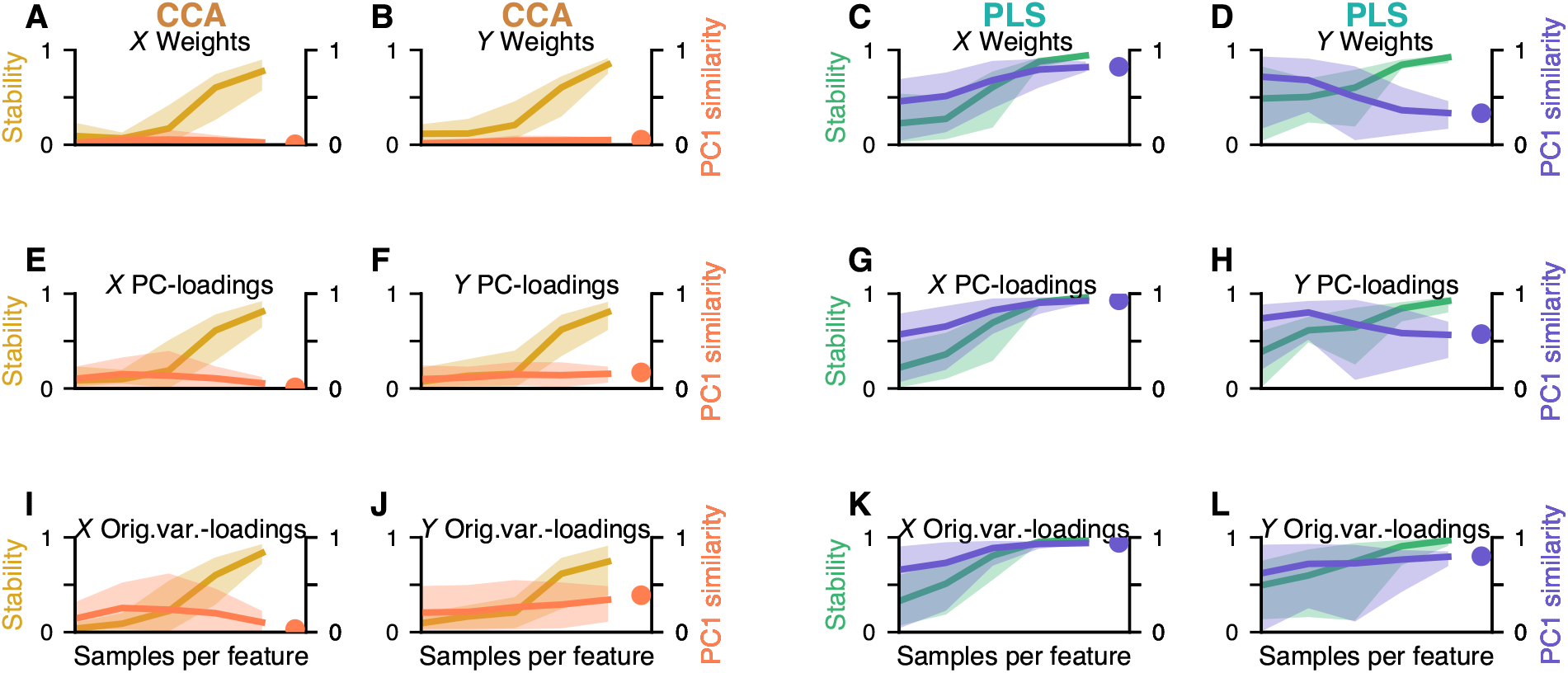
UKB fMRI CCA/PLS results for *X* and *Y* separately.

**Figure S12.**
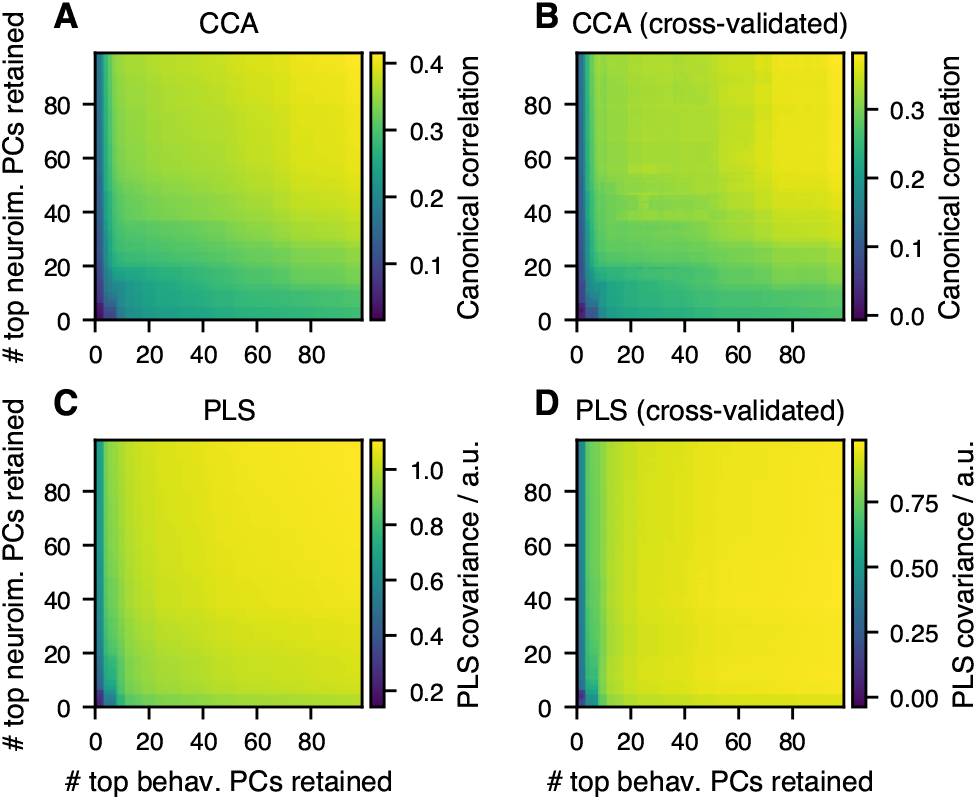
CCA and PLS association strength in UKB depending on retained number of principal components. **A)** In-sample and **B)** cross-validated association strength for CCA, measured as between-set correlation. **C)** In-sample and **D)** cross-validated association strength for PLS, measured as between-set covariance.

**Figure S13.**
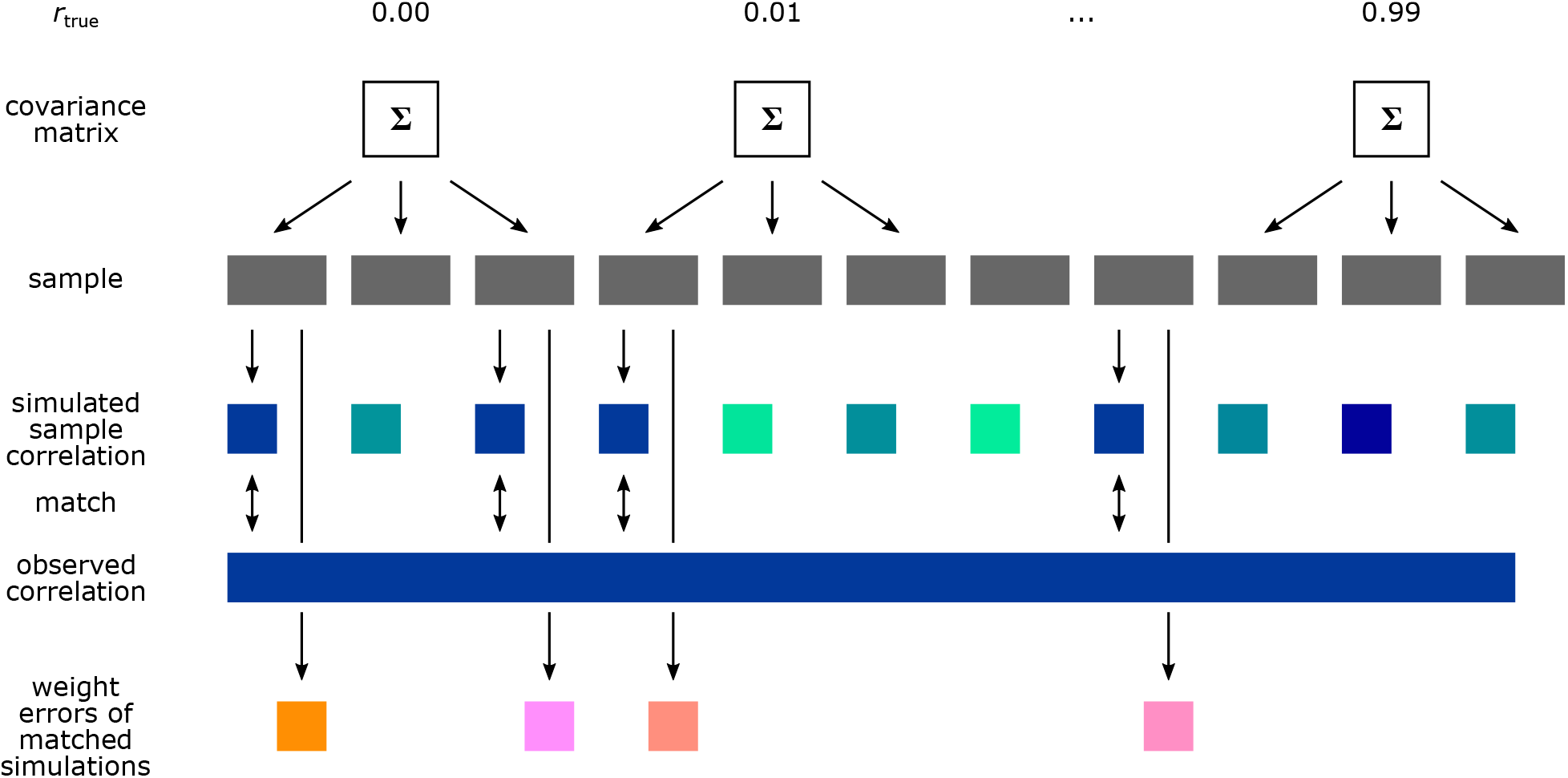
Schematic for estimating weight errors for published CCA results. For each CCA from the literature in our database, synthetic data for CCA is generated with matching number of samples and features. Separate datasets are generated for assumed ground-truth between-set correlations *r*_true_ varying between 0 and 0.99. In each generated dataset the canonical correlation is estimated and if it is close to the value in the reported CCA, the weight error for the synthetic dataset is recorded. The distribution of recorded weight errors across assumed ground-truth correlations and repetitions of the whole process is shown in Fig. 6B and its mean in Fig. 6A.

**Figure S14.**
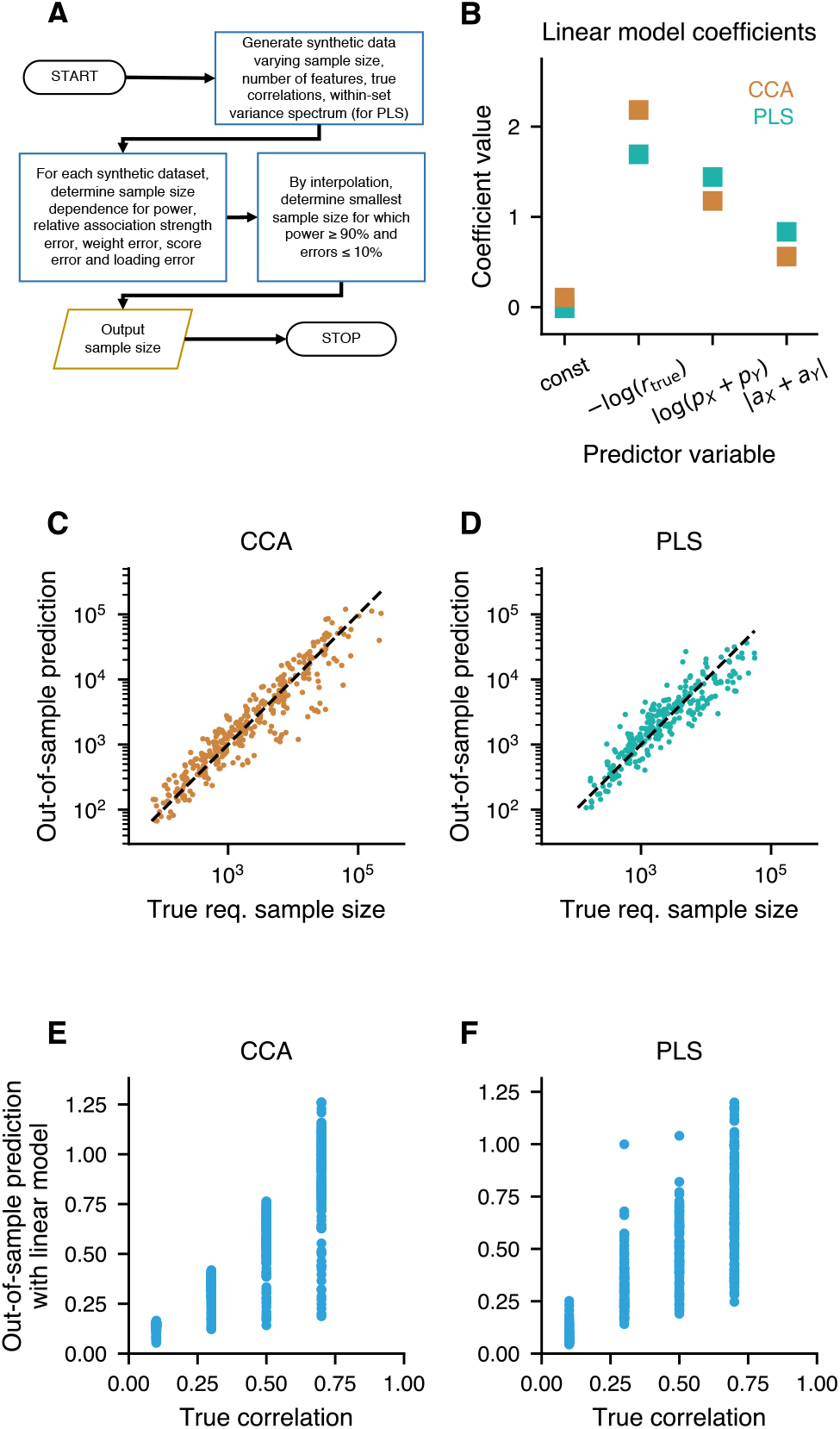
Sample-size calculator. **A)** Algorithm for sample-size calculation. Sample sizes can, in principle, be calculated directly with GEMMR, as shown in Fig. 6C-D. However, this is computationally expensive. To quickly obtain sample-size estimates, we developed the algorithm illustrated here. **B-G)** Especially for low assumed ground-truth correlations and a high number of features it is computationally expensive to estimate the required number of samples by generating synthetic datasets and searching the sample size such that error bounds are satisfied. To abbreviate this process we pre-calculate required sample sizes using the generative model approach for certain parameter values, fit a linear model to log(*n*_required_) and then use it to quickly interpolate for parameter values not in the pre-calculated database. Predictors for the linear model are *−* log(*r*_true_), log(*p_X_* + *p_Y_*) and, for PLS only, *|a_X_* + *a_Y_ |*, where *r*_true_ indicates the true between-set correlation, *p_X_* and *p_Y_* are the number of features in datasets *X* and *Y*, respectively, and *a_X_* and *a_Y_* are the power-law decay constants for the within-set principal component spectrum, respectively. Shown here are linear model estimates for the required sample size based on the combined criterion, i. e. the sample sizes required to obtain 90 % power and at most 10 % error for the between-set association strength, weight, score and loading error. **B)** Linear model coefficients for CCA and PLS. **C-D)** The pre-calculated database was split in half where one half corresponded to true between-set correlations of *r*_true_ = 0.1 and 0.3, the other to *r*_true_ = 0.5 and 0.7. The linear model was re-estimated separately for each half, and used to predict the other half. We obtained good predictions for CCA (**C**) and PLS (**D**). **E, F)** Solving the linear model for *r*_true_, we aim to predict correlations. We train the model using either simulation outcomes for *r*_true_ *∈ {*0.1, 0.3*}}*, or *r*_true_ *∈ {*0.7, 0.9*}}* and testing the predictions on the remaining *r*_true_s. **E)** Good predictions can be obtained in this way for CCA, **F)** but not for PLS.

**Figure S15.**
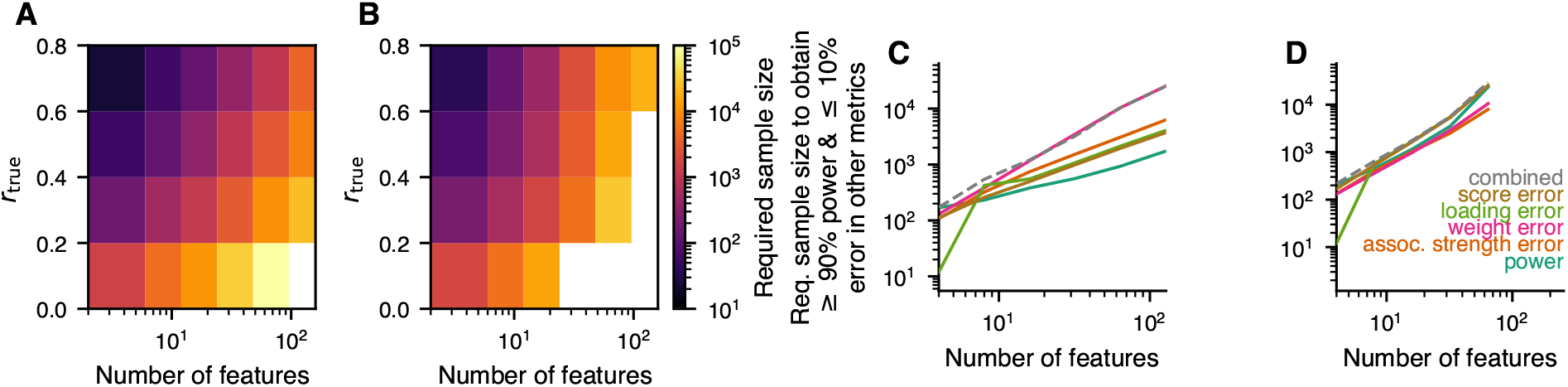
Parameter dependencies of required sample sizes. **A-B)** Required sample sizes based on the combined criterion increase with number of features and for low true between-set correlations *r*_true_. Due to computational expense values for some parameter sets were not available (white). **C-D)** Scaling of sample-size dependence on number of features, shown here for *r*_true_ = 0.3, for different metrics.

**Figure S16.**
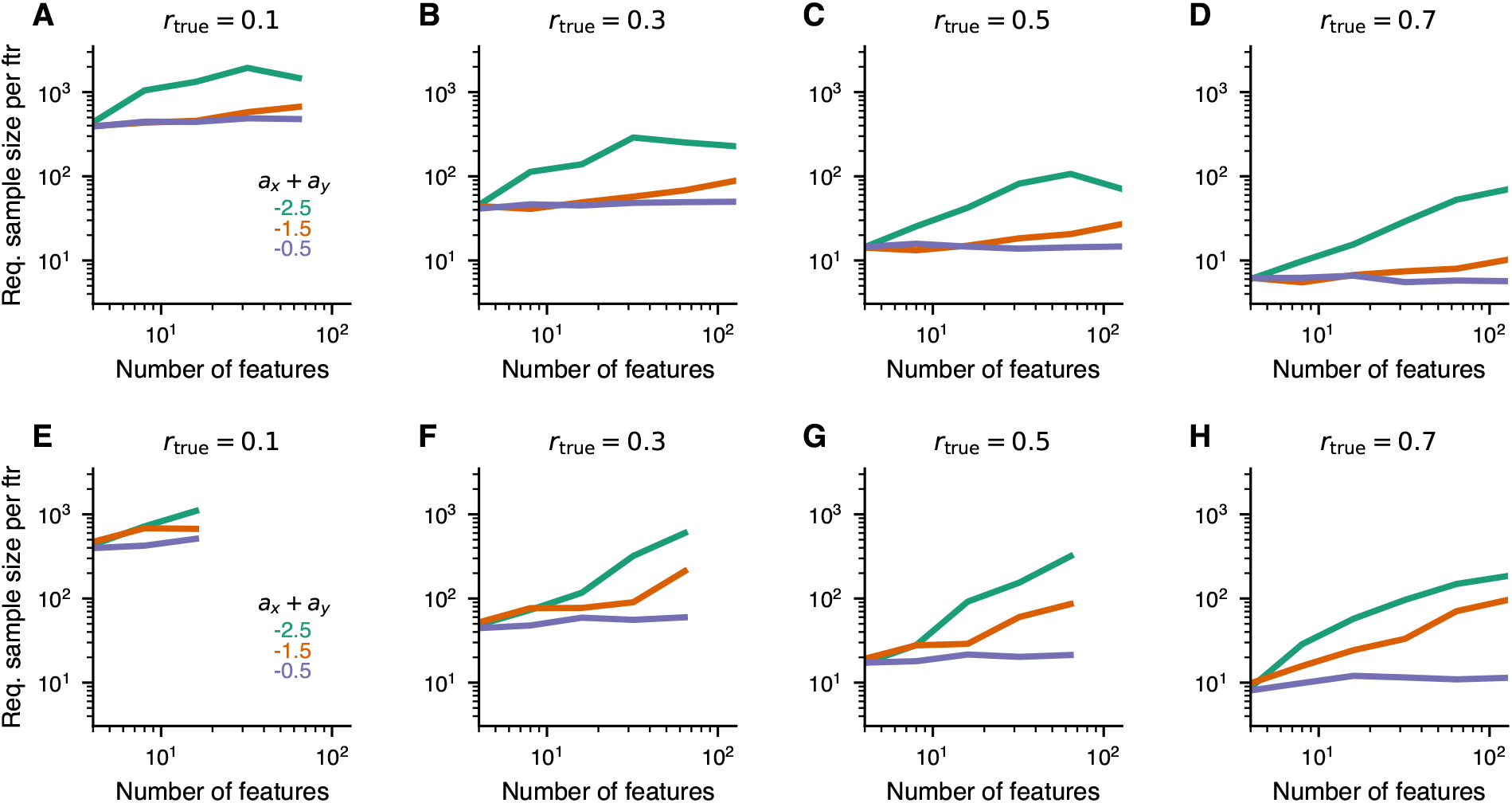
Sample-size dependence on within-set variances. **A-D)** CCA, **E-H)** PLS. Simulated parameter sets were averaged across subsets having indicated values for the between-set correlation *r*_true_ and for *a_X_* + *a_Y_* (the sum of within-set power-law decay constants) *±*0.5. The closer *a_X_* + *a_Y_* was to 0 (i. e. the “whiter” the data) the fewer samples were required.

**Figure S17.**
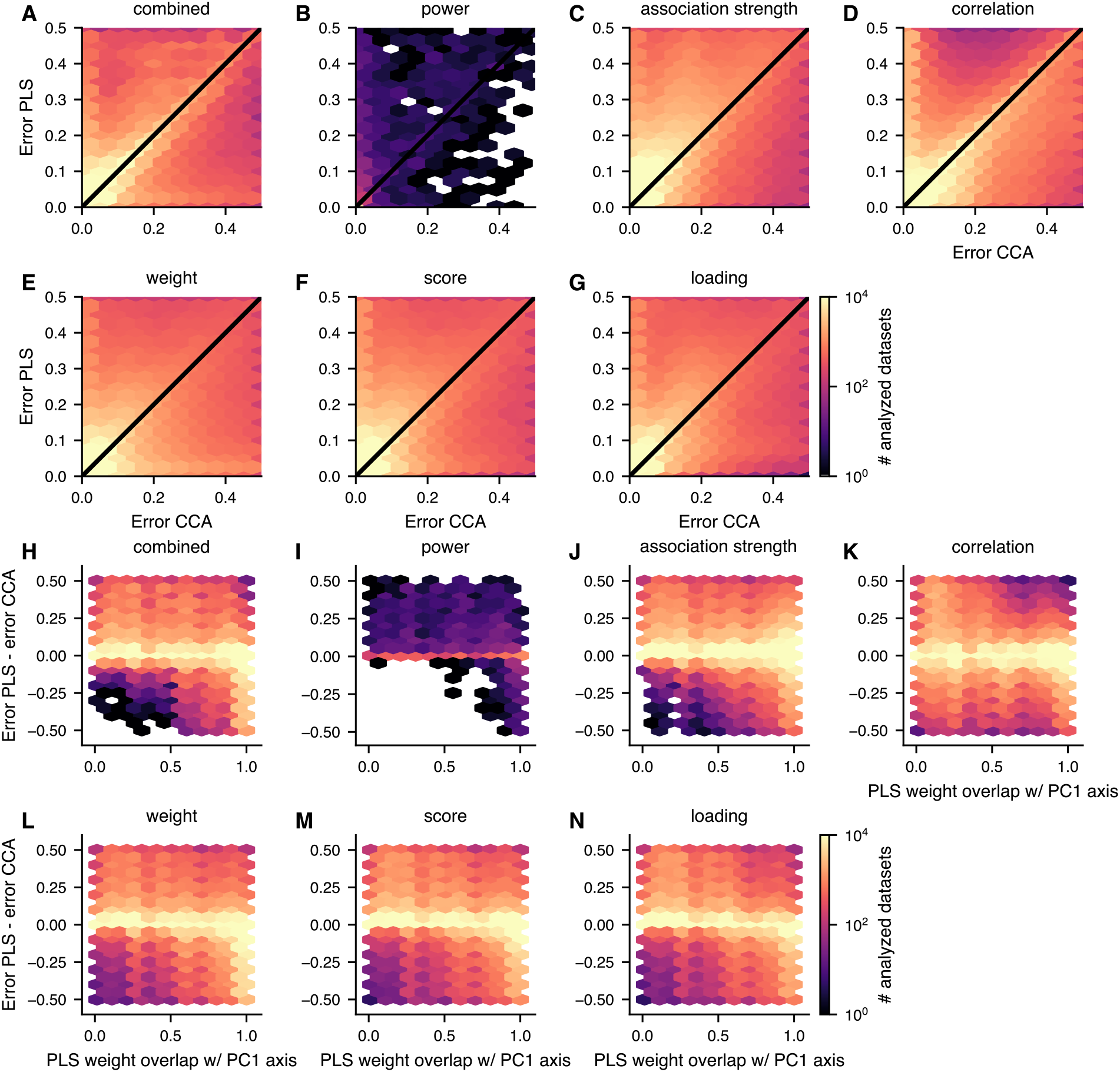
Required sample size for CCA vs PLS. We instantiated joint covariance matrices (assuming 1 between-set association mode), drew samples from the associated normal distributions, and analyzed the resulting datasets with both CCA and PLS. The CCA and PLS estimations were then compared, respectively, to the true CCA and PLS solutions, which were derived from the joint covariance matrices. Panels **A-G)** show for various error metrics how resulting deviations from the truth compare between CCA and PLS. PLS errors for a given dataset tend to be larger than CCA errors in many, but not all, datasets. **H-N)** For various error metrics, when PLS has a smaller error than CCA, this tends to happen preferentially when the true PLS weight overlaps strongly with the PC1 axis. Datasets were included in these analyses if the CCA or PLS error were below 0.5.

**Figure S18.**
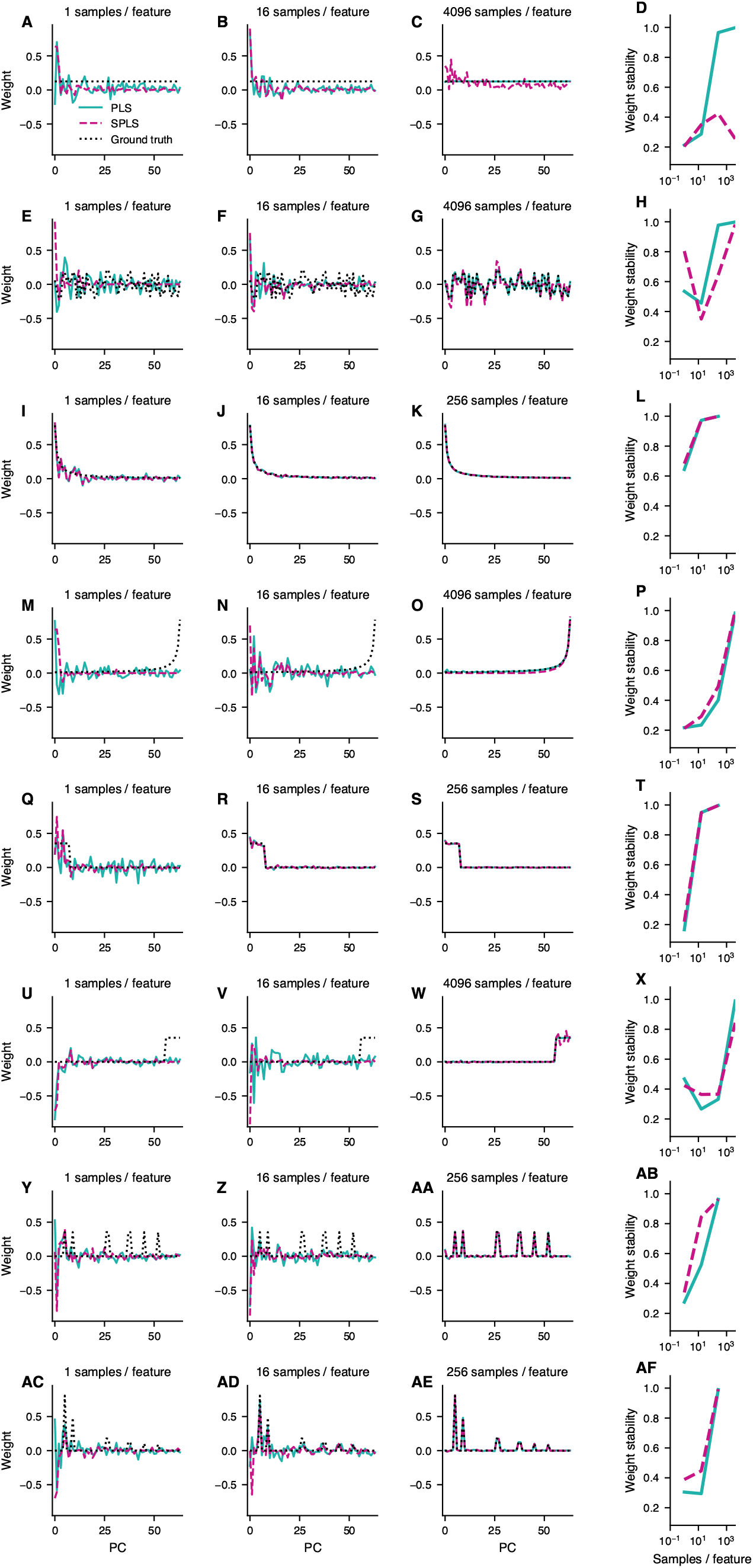
Comparison of PLS and sparse CCA (PLS). We have compared the capacity of PLS and sparse PLS [Witten et al., 2009] (this method is also referred to as “sparse CCA”, see Discussion) to recover different kinds of weight profiles (cf. different rows in figure to the left). For all except the first row, PLS and sparse PLS estimated the correct weight profile provided the sample size was large enough. In the first row, where a flat weight was assumed, sparse PLS failed, although it is unclear if even larger sample sizes would allow sparse PLS to succeed. Note that, for small sample sizes, both PLS and sparse PLS put strong weight on the first principal component, irrespective of the true population weight. Note also, that weight stability (last column) was, overall, similar for PLS and sparse PLS: sparse PLS had a slight advantage for sparse population weights and small sample sizes (last 2 rows). To produce these data, we assumed the same population weight for X and Y, a population between-set correlation of 0.3 and a within-set principal component variance spectrum decaying with constant -1 (for both X and Y). We ran 4 repetitions for each parameter combination and calculated stability as the pairwise cosine similarity between all possible pairs among the 4 repetitions. Altogether, in the situations show-cased here, sparse PLS did not do better than PLS.

## Supporting Information

### Canonical Correlation Analysis (CCA) and Partial Least Squares (PLS)

We assume that we have two datasets in the form of data matrices *X* and *Y*, both of which have *n* rows representing samples, and, respectively, *p_X_* and *p_Y_*columns representing measured features (or variables). Throughout we also assume that all columns of *X* and *Y* have mean 0. If both datasets consisted of only a single variable, we could measure their association by calculating their covariance or correlation. On the other hand, if one or both consist of more than one variable, pairwise between-set associations can be obtained but the possibly huge number of pairs results in a loss of statistical sensitivity and a difficulty to concisely interpret a potentially large number of significant associations [1]. To circumvent these problems, canonical correlation analysis (CCA) and partial least squares (PLS) estimate associations between weighted composites of the original data variables and find those weights that maximize the association strength.

### Terminology

Given a data matrix, e. g. *X*, composite variables or *scores t⃗_X_* (a vector of the same size as the number of samples, *n*) are formed by projection of *X* onto a *weight* vector *w⃗_X_* (of same size as the number of variables in *X*, *p_X_*), see Fig. 1A:

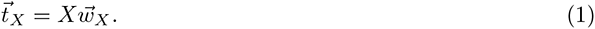

*Loadings ℓ⃗_XX_* (of same size as the number of variables in *X*, *p_X_*) characterize these composite variables by measuring their similarities with each of the original data variables in *X* (Fig. S18A-B)

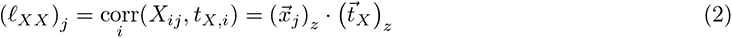

where corr means Pearson correlation, *x⃗_j_* is the *j*-th column of *X*, and the subscript *z* represents *z*-scoring across samples (i. e. subtraction of the mean and subsequent division by the standard deviation across samples). The complete loading vector is then

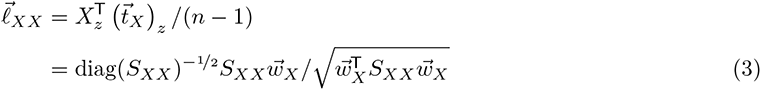

where *S_XX_* is the sample covariance matrix for *X*. Similarly, cross-loadings can be defined as

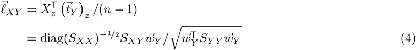

where *S_Y Y_* and *S_XY_*are, respectively, the sample covariance matrix for *Y* and the sample cross-covariance matrix between *X* and *Y*.

We note that alternative terminologies exist. CCA/PLS “scores” (as described above) could also be called “variates”, “weights” (as described above) could also be called “vectors”, and “loadings” (as described above) could also be called “parameters”. For CCA, the correlation between the score vectors, i. e. the “between-set correlations” or “inter-set correlations” are also called “canonical correlations”.

### Partial Least Squares

Partial Least Squares (PLS) finds the maximal covariance achievable between weighted linear combinations of features from two data matrices *X* and *Y* [2]:

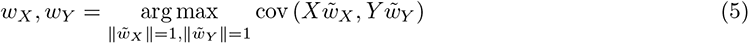

The solution is based on the between-set covariance matrix Σ*_XY_* which can be estimated from data via its sampled version 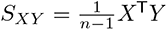. Performing a singular value decomposition yields

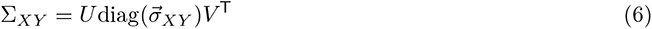

such that the optimal weights are given by the first columns of *U* and *V*, and the maximal covariance max

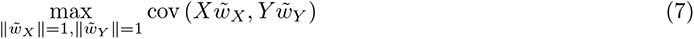

by the first singular value *σ_XY,_*_1_ [3, 4].

Multiple modes of association can be estimated in this way: beyond only the first column, every pair of corresponding columns in *U* and *V* provides another mode such that cov(*XCu_i_, Y Cv_i_*) (for 1 *≤ i ≤* min(*p_X_, p_Y_*)) is maximal given that the covariance of lower modes (those with indices *< i*) has already been accounted for. There are a number of different algorithms for PLS that differ conceptually in how these higher modes are estimated [2, 3]. The one presented above (sometimes called “partial least squares correlation” or PLS-SVD) was chosen for its similarity to canonical correlation analysis (see below). Another notable PLS algorithm is “PLS regression” which, in contrast to the above flavor, is asymmetrical in its handling of *X* and *Y* in that it estimates weighted composites (scores) for *X* and re-uses these as predictors for *Y* [2].

### Canonical Correlation Analysis

Canonical Correlation Analysis (CCA) [5], as a multivariate extension of Pearson’s correlation, finds maximal correlations between weighted linear combinations of variables from *X* and *Y* :

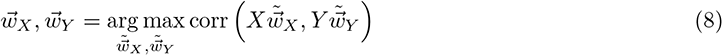

Note that corr 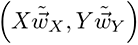 is independent of the scaling of 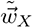 and 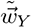. I. e. if *w⃗_X_* and *w⃗_Y_* are solutions of (8), then *c_X_w⃗_X_* and *c_Y_ w⃗_Y_*, where *c_X_ ∈* R and *c_Y_ ∈* R, are also solutions.

Also note that, as for PLS, several modes of association can be obtained with this framework by successively discounting the variance that has been explained by lower-order modes.

The maximal correlation in (8) is often called “canonical”.

The further analysis is then based on the “whitened” between-set covariance matrix

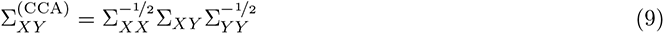

[6, 7]. A singular value decomposition of Σ^(CCA)^ is performed, yielding

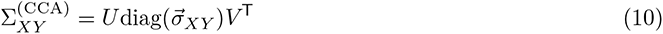

and the singular values *σ⃗_XY_*turn out to be the canonical correlations from (8), i. e. the maximal achievable correlations between a weighted linear combination of variables in *X* on the one hand, and a weighted linear combination of variables in *Y* on the other hand. The corresponding weights are given by

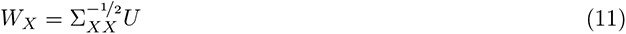

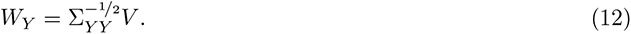

The use of the “whitened” between-set covariance matrix in CCA leads to an invariance property between datasets. To see this, let *X*_w_, *Y*_w_ be whitened data matrices, i. e. 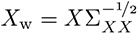 and 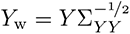 such that Σ*_X_*_w_ *_X_*_w_ = 1, Σ*_Y_*_w_ *_Y_*_w_ = 1. Then,

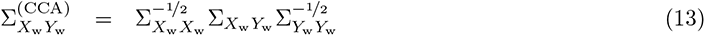

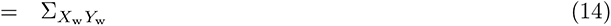

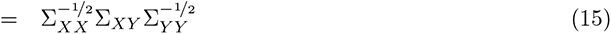

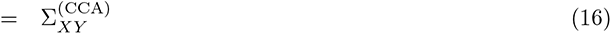

which is the same as for the original (non-whitened data). Consequently, canonical correlations for the original and whitened data are the same, given by the singular values of 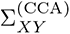, canonical weights for the whitened data are directly its singular vectors and canonical weights for the original (non-whitened) data differ only by a matrix 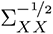 and 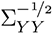 for *X* and *Y*, respectively (see (11)-(12)).

It can be shown that the invariance property is even more general [6]. Let *O*^(*X*)^ *∈* R*^pX ×pX^* and *O*^(^*^Y^* ^)^ *∈* R*^pY ×pY^* be non-singular and *d⃗*^(^*^X^*^)^ *∈* R*^pX^* and *d⃗*^(^*^Y^* ^)^ *∈* R*^pY^* be arbitrary vectors. Then *X̃* = *O*^(^*^X^*^)^*X* + *d*^(^*^X^*^)^ and *Ỹ* = *O*^(^*^Y^* ^)^*Y* + *d*^(^*^Y^* ^)^ have the same canonical correlations as *X* and *Y*, and the canonical vectors are related by

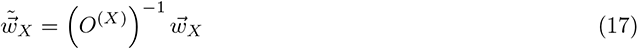

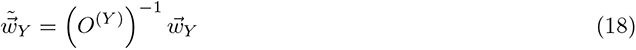

Thus, in particular, *z*-scored data *X_z_* = diag(*S_XX_*)*^−^*^1^*^/^*^2^*X* and *Y_z_* = diag(*S_Y Y_*)*^−^*^1^*^/^*^2^*Y* as well as whitened data *X_w_* and *Y_w_* have the same canonical correlations as the original data *X* and *Y*. In CCA, *X*- and *Y* -weights are related by [8]

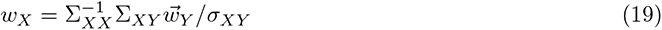

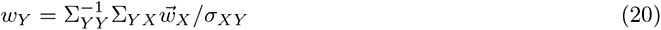

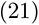

Replacing sample with population covariance matrices in (3) and (4), we thus also see that loadings and cross-loadings are collinear

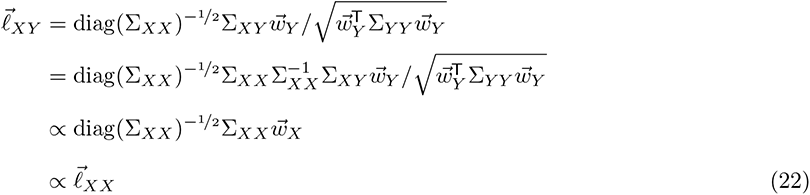

### Overestimation of association strength

Let Σ*_XY_* be a population cross-covariance matrix with singular value decomposition

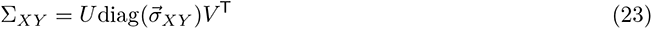

and let *_u*_1_, *_v*_1_ and *σ*_1_ be, respectively, the first columns of *U*, *V* and the first entry in *σ⃗_XY_*. In PLS, *_u*_1_, and *_v*_1_ are the weight vectors of the first mode and *σ*_1_ is the corresponding association strength.

The sample covariance matrix 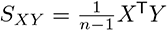 is an unbiased estimator for Σ*_XY_*, i. e. E[*S_XY_*] = Σ*_XY_*.

Therefore,

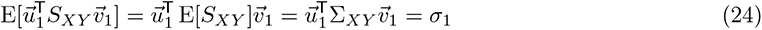

i. e. if the true (but unknown) weights were applied to a given dataset (between-set covariance matrix) the association strength of the resulting scores would, on average, match the true association strength. However, by definition, PLS selects those weight vectors that maximize the association strength between resulting scores. If 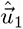 and 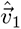 are those optimal weights for a given dataset, then

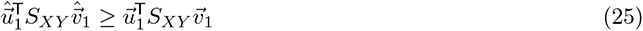

and consequently also

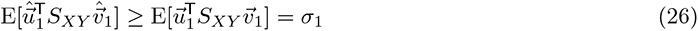

i. e. the association strength is overestimated.

### PC-1 similarity

We show here, for the special case that the between-set covariance matrix *S_XY_* has rank 1, that PLS weights are more similar to the first princpial component than CCA weights.

To describe the data we choose, without loss of information, a convenient coordinate system. Specifically, we assume that both *X* and *Y* data are expressed in their respective principal component coordinate system. Now, as said, we consider here the special case that *S_XY_* has rank 1, such that its singular value decomposition is

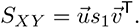

In this equation, *u⃗*_PLS_ *≡ u⃗* and *v⃗*_PLS_ *≡ v⃗* are, respectively, the *X* and *Y* PLS weight vectors.

To obtain the CCA solution, we consider 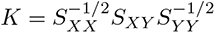. Then, we note that

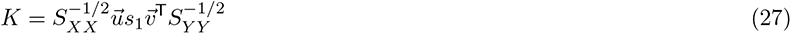

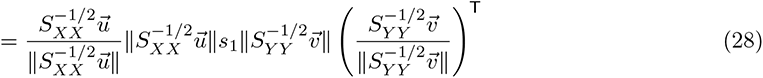

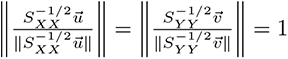, the right-hand side constitues the singular value decomposition of *K*. Thus 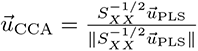 and 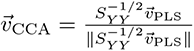 are the *X* and *Y* CCA weight vectors, respectively.

As we have assumed data are expressed in the principal component coordinate system, the cosine similarity of *u⃗*_PLS_ (analogously *v⃗*_PLS_) with the first principal component is simply the first coordinate of *u⃗*_PLS_, (*u⃗*_PLS_)_1_, which we denote *c*_PLS_. Likewise, the cosine similarity of *u⃗*_CCA_ with the first principal component is given by its first coordinate, denoted *c*_CCA_. To calculate that, let *λ*_1_ *≥ · · · ≥ λ_p_*denote the principal component variances for *X* and note that

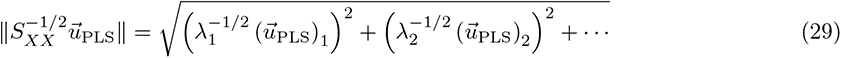

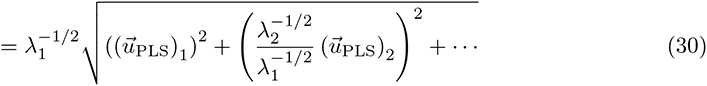

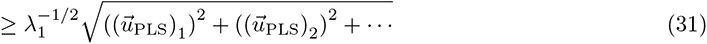

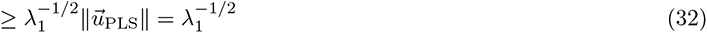

Then,

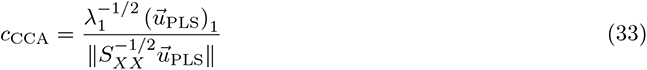

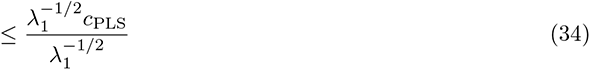

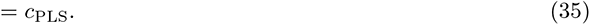

Thus, at least in the case of rank 1 between-set covariance matrices, PLS weight vectors have a stronger cosine similarity with PC1 than CCA weight vectors.

### Sparse CCA

Multiple sparse CCA and PLS methods exist [9–12]. Here, we use *penalized matrix decomposition* (PMD) [11], which has found widespread application, see e.g. [13–18]. Briefly, the PMD algorithm repeats the following steps until convergence [11]

- *u⃗ ←* arg max*_u⃗_ u*^T^*X*^T^*Yu⃗* subject to ||*u⃗*||_1_ *≤ c*_1_ and ||*u⃗*||_2_ *≤* 1
- *v⃗ ←* arg max*_v⃗_ u*^T^*X*^T^*Yv⃗* subject to ||*v⃗*||_1_ *≤ c*_2_ and ||*v⃗*||_2_ *≤* 1

to maximize *u⃗*^T^*X*^T^*Yv⃗*. If *X*^T^*X ≈* 1 and ||*u⃗*||_2_ = 1, then 1 = *u⃗*_2_ *≈ ||Xu||*_2_ and analogously for *Y*. Consequently, 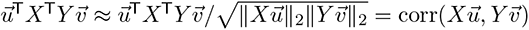. Note that the approximation *X*^T^*X ≈* 1 (together with *Y* ^T^*Y ≈* 1) makes this sparse “CCA” variant identical to sparse PLS [18, 19].

### Implementation and sparsity parameter selection

We implemented a Python wrapper for the R-package PMA [20] which we used with default parameters. Sparsity parameters were estimated separately for each dataset subjected to sparse CCA via 5-fold cross-validation [11, 21]: for *X* and *Y* we used 5 different candidate sparsity parameters (0.2, 0.4, 0.6, 0.8 and 1 where smaller values mean more sparsity and 1 corresponds to no sparsity) for a total of 25 parameter pairs. For each candidate parameter pair sparse CCA was estimated with 80 % of the data, the resulting weights applied to the remaining 20 % of the data to obtain test scores, the Pearson correlation calculated between the test scores and averaged across the 5 folds. The pair of sparsity parameters for which the test-correlation averaged across folds was maximal, was then selected and sparse CCA re-estimated on the whole data with these parameters.

